# Biomolecular condensation of cMLCK enables myosin motor phosphorylation in the heart

**DOI:** 10.64898/2026.05.21.726954

**Authors:** Kyler J. Carl, Austin Wellette-Hunsucker, Ivanka R Sevrieva, Kenneth S Campbell, Xu Fu, Elisabeth Ehler, Thomas Kampourakis

## Abstract

The heart needs to adapt its output to the metabolic demands of the organism. Phosphorylation of the myosin motors by cardiac myosin light chain kinase (cMLCK) increases heart muscle contractile function, yet its regulation and mechanism of action have remained unclear. Here, we show that cMLCK undergoes liquid-liquid phase separation and forms biomolecular condensates associated with the sarcoplasmic reticulum of cardiac muscle cells. Condensates selectively enrich enzymatic cofactors and substrates, which increases the catalytic activity of cMLCK. Our study reveals that cMLCK is fine-tuned to work in the molecular environment of condensates, enabling physiologically relevant levels of cardiac myosin motor phosphorylation. These findings establish a condensate-based mechanism for the spatial and temporal regulation of cardiac thick filament contractile function.

## Introduction

In addition to the well-studied classical Ca^2+^-dependent thin filament regulatory pathway of muscle contraction, activation of the myosin motors proteins in the thick filaments themselves has emerged as a second regulatory step that controls cardiac muscle contractile function (*1, 2*). Similar to the thin filaments, myosin motors in the thick filaments are believed to exist in both a diastolic OFF and systolic ON state, and the rate of transition between those states are likely rate-limiting for systolic force development and diastolic relaxation of the heart (Fig. 1A).

**Figure 1.**
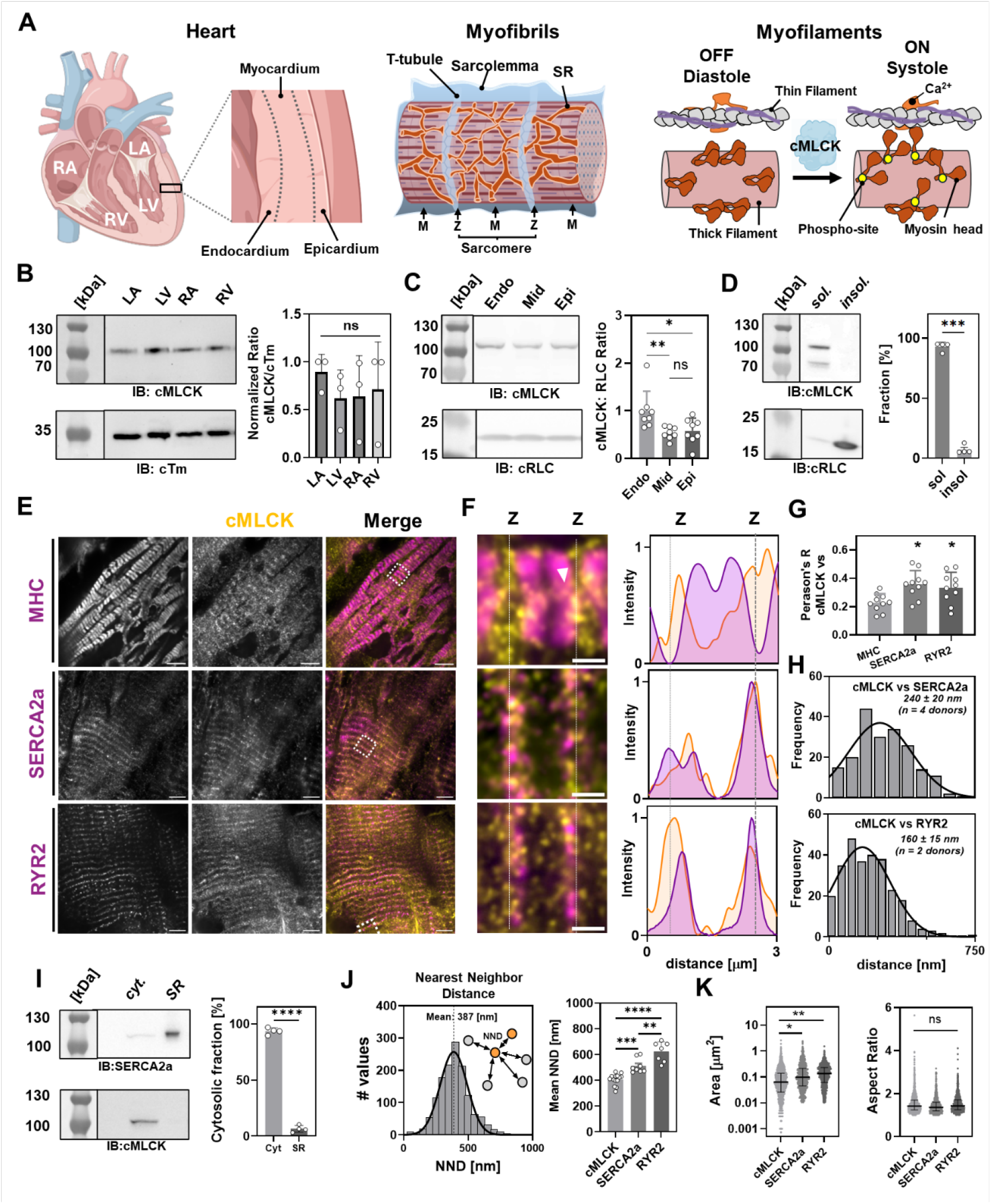
cMLCK is localized in cluster on the sarcoplasmic reticulum. (A) Structural organization of the heart at the organ, cellular and myofilament level. (B) Immuno-blot analysis of cMLCK expression in the left and right ventricles and atria of the human heart (mean ± s.d. for N=3). (C) Expression gradient of cMLCK through the ventricular wall (mean ± s.d. for N=8). (D) Fractionation of ventricular tissue into detergent-extracted soluble and insoluble fraction and analyzed by immune-blotting against both cMLCK and its substrate cRLC (mean ± s.d. for N=4). (E) Confocal SoRA super-resolution images of cryo-sections from human ventricular tissue immuno-stained against cMLCK, and counter-stained against myosin heavy chain (MHC, top), SERCA2a (middle) and ryanodine receptors (RYR2, bottom). Scale bars represent 5 μm. (F) Left: Expanded views of single sarcomeres shown in (E). Scale bars represent 1 μm. Right: Averaged longitudinal intensity profiles. (G) Co-localization analysis of cMLCK vs MHC, SERCA2a and RYR2 (pooled data from N=3, n=10). (H) Average distances between cMLCK, and SERCA2a (top) and RYR2 (bottom) clusters. (I) Distribution of cMLCK in the cytosolic (cyt) and sarcoplasmic reticulum (SR) fraction analyzed by immune-blotting against SERCA2a (top) and cMLCK (bottom) (N=4). (J) Nearest neighbor distribution of cMLCK, SERCA2a and RYR2 (N=3, n=7-14) (K) Size and shape distribution of cMLCK, SERCA2a and RYR2 clusters (pooled data from N=4). Data are shown as mean ± SD and statistical significance of differences were analyzed with a one-way ANOVA followed by Tukey’s or Dunn’s post-hoc test for parametric and non-parametric data sets, respectively. *p<0.05, **p<0.01, ****p<0.0001; ns – not significant.

Phosphorylation of the myosin motors controls cardiac output and performance by modulating both their availability for contraction and activity, and myosin motors are frequently found hypophosphorylated in human myocardium from patients with heart disease and heart failure (*3-6*). This underlines the functional significance of myosin phosphorylation for the normal performance of the heart. The primary enzyme responsible for phosphorylating the myosin motors in the heart is the cardiac isoform of myosin light chain kinase (cMLCK) (*7, 8*), which exhibits unique cardiac-specific features, including a low intrinsic activity and a large N-terminal domain (NTD) without a recognized function (*9*). The regulatory mechanisms underlying cMLCK localization and function have remained elusive and remain a barrier for the development of therapeutic interventions aimed at modulating myosin motor activity.

Biomolecular condensates, membrane-less organelles formed by liquid-liquid phase separation LLPS) of proteins and nucleic acids, have emerged as a fundamental mechanism for organizing biochemical reactions within cells (*10, 11*). LLPS is frequently driven by multivalent, low-affinity interactions involving intrinsically disordered regions (IDRs) and enables the spatial concentration of enzymes, substrates, and cofactors in ways that can profoundly alter reaction kinetics (*12-14*). While LLPS has been characterized extensively in the context of transcription regulation, RNA processing, stress responses and pathologies such as cancer and neurodegenerative diseases (*15-18*), its role in regulating heart muscle contractile function has not been explored.

In the current study, we demonstrate that cMLCK undergoes LLPS driven by its intrinsically disordered NTD, forming biomolecular condensates on the surface of the sarcoplasmic reticulum (SR) in human cardiomyocytes. We show that the condensate microenvironment is required for physiologically relevant cMLCK activation and cRLC phosphorylation, revealing a previously unrecognized mechanism for the regulation of cardiac thick filament contractile function.

## Results

### cMLCK localizes in c;lusters to the sarcoplasmic reticulum in human cardiomyocytes

To investigate the role of cMLCK in regulating the contractile performance of the human heart we analyzed its expression profile in human donor heart samples. Immuno-blot analysis using an antibody against the cardiac-specific N-terminal domain showed no difference in cMLCK expression across the four chambers of the heart (fig. S1 and Fig. 1B). However, analysis of the sub-myocardial layers revealed an expression gradient of cMLCK with the highest levels in the endocardium followed by the mid- and epicardial layers (Fig. 1C), consistent with previous reports of a potential transmural gradient of myosin RLC phosphorylation in the ventricular wall (*19*). This gradient has been proposed to facilitate cardiac torsion dynamics and ejection of blood by counteracting the greater wall stress experienced by the endocardial layers. We used quantitative immuno-blotting to estimate the cMLCK concentration in human cardiomyocytes from the mid-myocardial section to about 3.5 μmol L^-1^ (fig. S1B), in excellent agreement with previous measurements in rodent myocardium (*20*). Moreover, Phostag™-immunoblot analysis showed that cMLCK is phosphorylated to about 1.1 ± 0.1 mol Pi mol cMLCK^-1^ in-vivo (fig. S1C). In contrast to its high substrate specificity towards cardiac myosin cRLC, fractionation of ventricular tissue into myofilament and cytosolic fraction showed that <5% of cMLCK is bound to the myofilaments in human donor samples (Fig. 1D).

Sub-cellular localization is critical for protein kinase function by regulating both catalytic activity and proximity to its substrates. We therefore stained cryo-sections of intact left ventricular tissue isolated from human donors against cMLCK and its substrate cardiac myosin, and determined their localization using super-resolution imaging (Fig. 1E). In agreement with previous reports, cMLCK is mainly localized close to the Z-disc of the myofilaments with only little co-localization with its substrate cardiac myosin in the sarcomeric A-band. Rather than the continuous staining expected for a Z-disc associated protein, cMLCK localizes to distinct clusters reminiscent of a signal for junctional and longitudinal sarcoplasmic reticulum (SR) (Fig. 1F). We confirmed this interpretation by staining cryosections against both cMLCK, and the Ca^2+^ pump SERCA2a or Ca^2+^ release channel RYR2 in the SR, which showed a significantly higher degree of co-localization as indicated by Pearson’s correlation analysis (Fig. 1G). The calculated distance distribution between the centroids of both SERCA2a and RYR2 with the cMLCK clusters shows an average distance of about 150 nm and 250 nm, respectively (Fig. 1H). The proximity of cMLCK clusters to Ca^2+^ handling proteins on the SR likely enables tight control over their Ca^2+^/calmodulin-dependent activity. However, cMLCK does not co-purify with SR vesicles during tissue fractionation, suggesting that cMLCK clusters are not tightly bound to SR membranes and likely highly dynamic (Fig. 1I). In agreement, PIP strip assay showed no significant interaction between isolated cMLCK and SR membrane lipids (fig. S2).

The distribution of the distance between two nearest cMLCK puncta on the SR is Gaussian-like with a mean value of about 400 nm, suggesting a non-random distribution of cMLCK puncta on the SR membrane (Fig. 1J). In contrast, the average nearest neighbour distribution is significantly larger for both SERCA2a and RYR2. The average cMLCK cluster size can be estimated based on these values to about 1000 molecules (see Supplementary Text). The cMLCK clusters follow a log-normal size distribution with a median area of about 0.05 μm^2^ (equivalent of a circle of 130 nm radius), which is smaller than SERCA2a and RYR2 clusters and show an approximately circular shape as indicated by an average aspect ratio close to 1 (Fig. 1K).

Collectively, these data indicate that cMLCK is not constitutively associated with the myofilaments in human donor myocardium, but is instead organized into discrete, SR-associated clusters near calcium-handling proteins SERCA2a and RYR2.

### cMLCK forms biomolecular condensates via liquid-liquid phase separation

The clustering of cMLCK into distinct puncta suggests the formation of biomolecular condensates on the surface of the membrane of the SR. Membranes reduce protein diffusion and locally increase protein concentrations, which facilitates condensate formation via liquid-liquid phase separation (LLPS) (*21, 22*). LLPS is frequently driven by multivalent and dynamic interactions of intrinsically disordered regions of proteins. Bioinformatics analysis of cMLCK showed that its unique N-terminal domain (NTD) is largely disordered with a high propensity to drive LLPS, apart from a potential coiled-coil structure within the first 150 amino acids (Fig. 2A). We tested this idea by expressing either full length (FL) cMLCK, its isolated NTD or catalytic subunit (catSU) fused to eGFP in both primary neonatal rat cardiomyocytes (NRCs) and COS-1 cells (Fig. 2B). Both FL cMLCK and its NTD readily formed condensate-like structures in both cell types. We excluded effects due to protein fusion by comparing the size and shape distribution of cMLCK fused to either eGFP or FusionRed (fig. S3A). FusionRed has been shown to have minimal effect on protein condensate formation (*23*). Moreover, cell type did not affect the distribution of cMLCK condensate size or shape, suggesting that phase separation is an intrinsic property of cMLCK NTD (fig. S3B). Lack of colocalization with p62 (SQSTM1) further indicates that these structures are not autophagosomes (fig. S3C). Although both NTD and FL cMLCK form condensates of roughly equal size distribution, the NTD forms more abundant condensates that exhibit a more dynamic shape distribution (Fig. 2C), suggesting that LLPS of cMLCK is driven by its NTD but modulated by its catalytic subunit.

**Figure 2.**
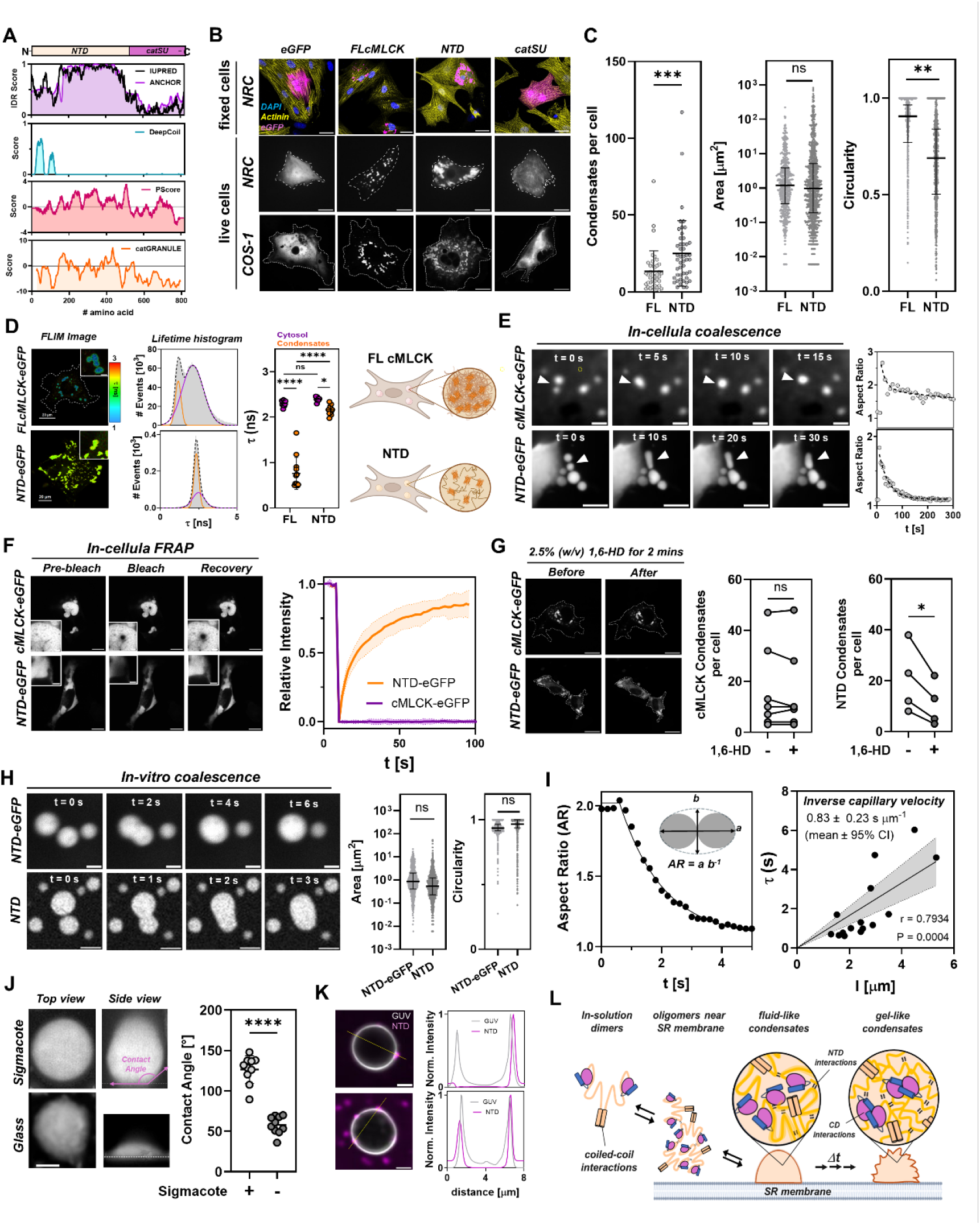
cMLCK forms biomolecular condensates via liquid-liquid phase separation. (A) Bioinformatics analysis of cMLCK. (B) Confocal microscopy images of COS-1 cells and neonatal rat cardiomyocytes (NRCs) expressing eGFP alone or fused to cMLCK constructs (FL, full length; NTD, N-terminal domain; catSU, catalytic subunit). Scale bars represent 10 μm. (C) Comparison of cMLCK-eGFP and NTD-eGFP condensate abundance, size and shape distribution (pooled data from N=4). (D) Fluorescence life-time imaging of FL cMLCK- and NTD-eGFP condensates in COS-1 cells (N=3, n=9-13). (E) Live cell imaging of coalescence of cMLCK-eGFP (top) and NTD-eGFP (bottom) condensate.

We probed the chemical environment of FL cMLCK and NTD condensates using fluorescence lifetime imaging (FLIM) (Fig. 2D). eGFP conjugated to either FL cMLCK or NTD localized in the cytosol exhibits a fluorescence lifetime (τ) of about 2.4 ns, in good agreement with measurements on isolated eGFP (*24*). However, the eGFP fluorescent lifetime was strongly reduced in FL cMLCK condensates (about 1 ns), indicative of a highly crowded and viscous environment with high refractive index (*25*). In contrast, eGFP fluorescence lifetime was only modestly reduced in condensates formed by the NTD (about 2.2 ns), suggesting a less dense micro-environment.

Coalescence time courses are shown on the right. (F) In-cellula fluorescence recovery after photobleaching experiments with cMLCK-eGFP and NTD-eGFP condensates (pooled data from N=3, n=3-8). (G) Effect of 1,6-Hexandiol treatment of cMLCK-eGFP and NTD-eGFP condensates (N=2-3, n=4-7). (H) Confocal images time-series of coalescence assay of condensates formed from bacterial expressed and purified isolated NTD either fused to eGFP (top) or mixed with 5% (mol mol^-1^) Alexa647 – NTD (bottom). (I) Determination of the inverse capillary velocity of condensates by linear regression of fusion time (τ) plotted against condensate size (l). (J) Effect of surface treatment on condensate wetting of glass surfaces. Contact angles (n=10). (K) Interaction of NTD condensates (magenta) with GUV (gray). Line scans are shown on the right. (L) Cartoon representation of the proposed mechanism of cMLCK condensate formation on the surface of the SR membrane. Data in (C) and (H) were analyzed using a nested t-test. Data in (D) were analyzed using a two-way ANOVA followed by Tukeys multiple comparison test. Data in (G) and (J) were analyzed using an unpaired student’s t-test.

Both FL cMLCK and NTD condensates coalesced in in-cellula experiments, however, with significantly different time-courses (Fig. 2E). Moreover, cMLCK condensates showed frequent arrested fusion and fission events (fig. S4A). NTD condensates showed a rapid recovery in fluorescence recovery after photobleaching (FRAP) experiments with a halftime of about 12 s (Fig. 2F), confirming their liquid-like internal structure. In contrast, cMLCK condensates showed only little recovery in FRAP. This suggests that condensates in the presence of the catalytic subunit are in a gel-like visco-elastic state, consistent with the FLIM measurements and arrested fusion events. NTD condensates readily dissolved in-cellula during a brief treatment with 1,6-Hexandiol (1,6-HD), which has been shown to disrupt the weak intermolecular interactions that stabilize fluid-like condensate architecture, whereas cMLCK condensates were largely resistant to 1,6-HD (Fig. 2G). Taken together, this suggests that in the presence of the catalytic subunit, cMLCK condensates undergo a liquid-to-gel transition. Prolonged culture of COS-1 cells shows that NTD condensates undergo a similar transition, as indicated by a reduced recovery in FRAP experiments, but at a significantly slower time-scale (fig. S4B).

To test whether condensate formation is an intrinsic property of the NTD and does not depend on the cellular environment or the presence of other factors, we purified both recombinantly expressed eGFP-tagged and unlabeled NTD, and tested for LLPS via confocal microscopy. Phase separation of the unlabeled NTD was monitored by addition of 5% (mol mol^-1^) of Alexa647-labeled NTD. Both proteins underwent spontaneous LLPS to form similar sized and well-defined liquid-like condensates that readily fused under in-vitro conditions (fig. S5A and Fig. 2H). LLPS and coalescence of NTD were also observed via phase contrast microscopy in the absence of fluorescent labels (fig. S5B). We measured the fusion-time as a function of condensates size to estimate an inverse capillary velocity of about 0.8 s μm^-1^ (Fig. 2I), suggesting a low ratio of viscosity to surface tension. Condensates also showed wetting of glass surfaces as expected for a liquid-like state with contact angles >120º, which could be reduced by increasing the hydrophobicity of the glass surface via silanization (Fig. 2J), suggesting that condensates readily interact with hydrophilic and charged surface. To test if NTD condensates can interact with the polar surface of lipid bilayers, we prepared giant unilamellar vesicles (GUV) from 1,2-dioleoyl-sn-glycero-3-phosphocholine (DOPC) mixed with 0.1% (mol mol^-1^) 1,2-dioleoyl-sn-glycero-3-phosphocholine labeled with ATTO-488 for visualization via confocal microscopy (Fig. 2K). The NTD formed small round condensates on the surface of the GUVs, reminiscent of the cMLCK clusters on the surface of the SR observed in human cryosections (Fig. 1E). Similar to the in-cellula experiments, isolated NTD condensates underwent a time-dependent liquid-to-gel like transition associated with the emergence of amyloid fibril-like structures as indicated by Thioflavin T staining (fig. S5C) and slow recovery in in-vitro FRAP experiments (fig. S5D).

To further investigate the mechanism of cMLCK condensate formation, FusionRed-labeled cMLCK was co-expressed with either eGFP-labeled catSU or NTD (fig. S6A). Both NTD and catSU were enriched in FL cMLCK condensates with the NTD showing a higher degree of co-localization. This suggests that both domains might form inter-molecular interactions that contribute to condensate architecture. In fact, both proteins undergo homo-dimerization and oligomerization in-vitro (fig. S6B), although this effect was significantly weaker for the catSU. The isolated NTD undergoes dimerization with high affinity (K_d_ of about 40 nmol L^-1^) (Fig. S6C), suggesting that the kinase exists as a soluble dimer. Deletion of the coiled-coil sequence completely abolished phase separation of the NTD, indicating that dimerization is a requirement for biomolecule condensate formation of cMLCK (fig. S6D).

Taken together, our results suggest a mechanism in which cMLCK dimers undergo rapid LLPS via dynamics interactions of their disordered region of the NTD into liquid-like condensates on the SR membrane. This is followed by internal crosslinking of cMLCK molecules via static interactions of both their coiled-coil and catalytic subunits domains, associated with a maturation into a gel-like visco-elastic network (Fig. 2K).

### Ca^2+^ activation recruits calmodulin into condensates and activates cMLCK

Previous studies have shown a strong Ca^2+^/calmodulin-dependent activity of cMLCK (*3*). To study the effect of calmodulin (CaM) activation of cMLCK in the context of biomolecular condensates, we co-expressed FL cMLCK-eGFP with mCherry-fused CaM in NRCs (Fig. 3A and fig. S7A). CaM was actively recruited into cMLCK condensates as shown by high degree of co-localization. Strikingly, however, co-expression had no effect on the number of cMLCK condensates per cell, or their size or shape distributions (Fig. 3B), suggesting that Ca^2+^/CaM activation of cMLCK does not release the kinase from its condensates.

**Figure 3.**
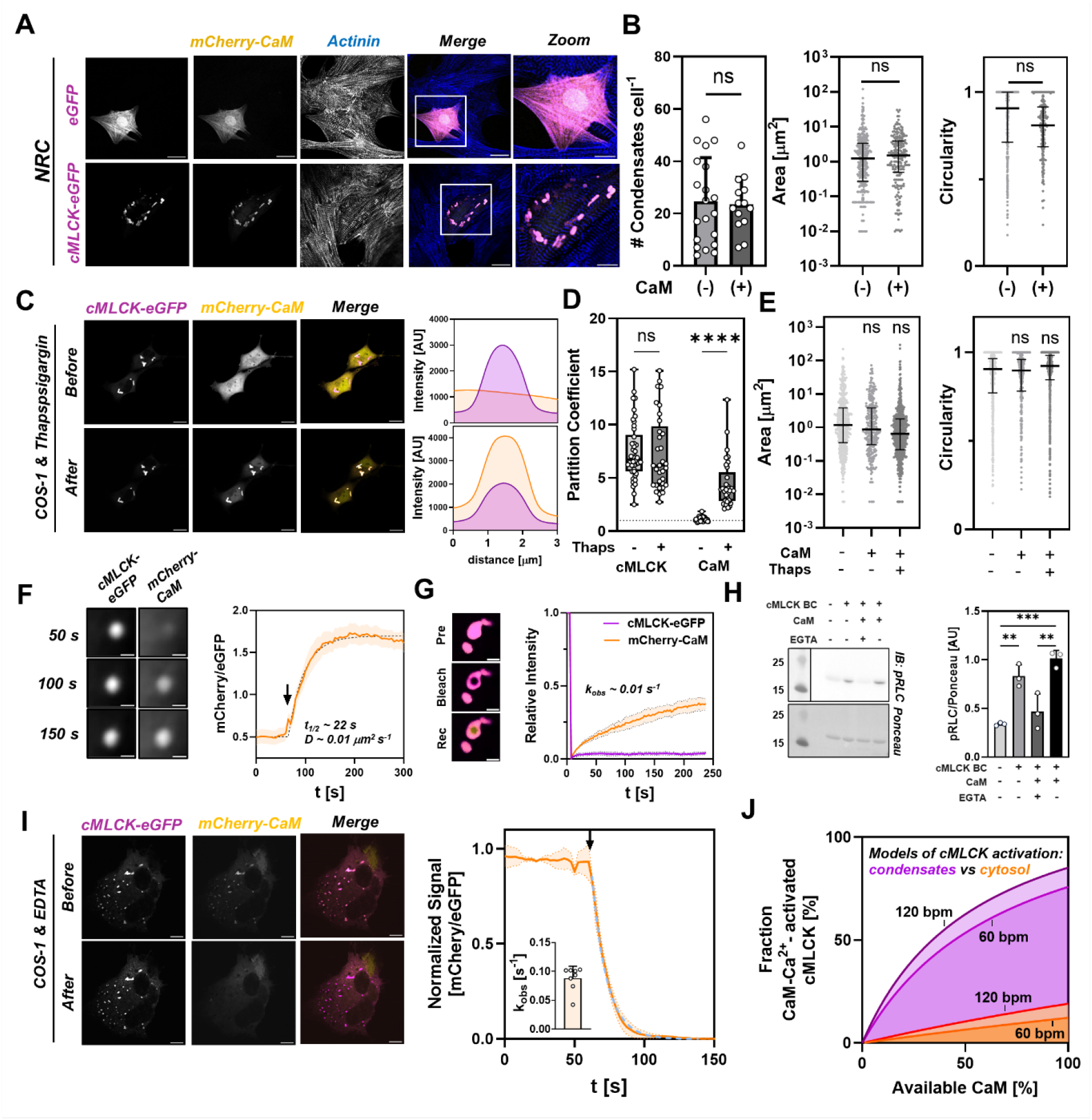
Calcium activation recruits calmodulin into condensates and activates cMLCK. (A) Co-expression of eGFP (top) or cMLCK-eGFP (bottom) with mCherry-CaM in NRCs. (B) Comparison of cMLCK-eGFP condensate abundance (left), and size (middle) and shape distribution (right) in the absence (-) and in the presence of mCherry-CaM (+) in NRCs (pooled data from N=2 repeats, n =20 cells). (C) Localization of cMLCK-eGFP with mCherry-CaM in COS-1 cells before and after treatment with Thapsigargin. Intensity profile of the line scans are shown on the right. (D) Partition coefficients of cMLCK-eGFP and mCherry-CaM in condensates before (-) and after Thapsigargin treatment (+) of COS-1 cells (pooled data from N=3, n=33-44 cells). (E) Size and shape distribution of cMLCK-eGFP condensates in the absence and in the presence of mCherry-CaM (CaM), and after treatment of cells with Thapsigargin (Thaps) (pooled data from N=3, n=33-44 cells). (F) Time-course of mCherry-CaM recruitment into cMLCK-eGFP condensates (N=3, n=8). (G) Dual-color FRAP of cMLCK-eGFP and mCherry-CaM in COS-1 cells treated with Thapsigargin (N=3, n=8). (H) In-vitro kinase assay with purified cMLCK-eGFP condensates (N=4 repeats). (I) Time-course of mCherry-CaM dissociation from cMLCK-eGFP condensates in saponin-treated COS-1 cells after treatment with EDTA (addition of EDTA is indicated by a black arrow) (N=3, n=8). (J) Model comparison of activation levels of cMLCK either localized to condensates (purple) vs freely diffusible in the cytosol (orange) for 60 bpm and 120 bpm. Data in panel (B) and (D) were analyzed using a nested unpaired, two-tailed t-test for either normal or lognormal distributed data sets. Data in panels (E) and (H) were analyzed with a nested one-way ANOVA followed by Tukey’s multiple comparison test.

NRCs are excitable cells and exhibit spontaneous calcium transients. To more closely study the recruitment of calmodulin into cMLCK condensates and kinase activation, we used the simpler COS-1 cell system (Fig. 3C). The basal cytosolic Ca^2+^ concentration in COS-1 cells is maintained at about 100 nmol L^-1^ (*26*), similar to the diastolic value observed in cardiomyocytes. Under these conditions, CaM is not recruited into cMLCK condensates but can freely percolate the condensate space as indicated by a partition coefficient ([condensate] [cytosol]^-1^) close to 1 (Fig. 3D). To study the effect of calcium activation we treated COS-1 cells with Thapsigargin, which led to a fast, reproducible and sustained increase in cytosolic Ca^2+^ with a halftime of about 8 s (fig. S7B). Thapsigargin treatment of COS-1 cells led to a recruitment of CaM into cMLCK condensates as indicted by a partition coefficient of about 5-10, which was linear related to the amount of cMLCK in the condensates, suggesting specific recruitment (fig. S7C). However, consistent with the observations in NRCs, calcium activation and CaM recruitment had no effect on the either the size or shape distribution of cMLCK condensates (Fig. 3E). We quantified the kinetics of CaM recruitment into condensates using time-resolved super-resolution imaging (Fig. 3F), showing a halftime of about 22 s and an estimated diffusion coefficient of about 0.01 μm^2^ s^-1^. This value is several orders of magnitude lower than what was observed for free diffusion of proteins in the cytosol (*27*), consistent with the highly crowded and visco-elastic micro-environment of the cMLCK condensates. We probed the internal dynamics of CaM in condensates using FRAP, which showed a rate of recovery of about 0.01 s^-1^, likely reflecting the OFF rate of CaM-Ca^2+^ from cMLCK in the condensate. We used Microscale Thermophoresis and tryptophan fluorescence to quantify the steady-state dissociation constant (K_d_) of calmodulin binding to cMLCK to about 1 μmol L^-1^, therefore giving an ON rate of about 10000 mol L^-1^ s^-1^ (fig. S8A). We next tested if cMLCK condensates exhibit catalytic activity by enriching condensates from COS-1 cells using differential centrifugation and using them in in-vitro kinase assays with recombinant cRLC as a substrate (fig. S8B and Fig, 3H). The purified condensates showed phosphorylation activity towards cRLC, which could be inhibited by EGTA. Adding an excess of CaM and Ca^2+^ restored cMLCK activity in condensates towards cRLC. These results strongly suggest that Ca^2^ activation has no effect on cMLCK condensate formation but rather leads to recruitment of CaM and activation the enzymatic activity of cMLCK inside the condensate environment.

We also tested for the release kinetics of CaM from cMLCK in condensates using saponin-permeabilized COS-1 cells, which allow rapid diffusion of small molecules into the cytosol but prevents protein leakage (Fig. 3I). Addition of excess EDTA to Ca^2+^-treated, permeabilized COS-1 cells shows a rapid decline in co-localization of mCherry-CaM with cMLCK-eGFP condensates, which is best described by a single exponential decay with a rate constant of about 0.1 s^-1^. This is about three times faster than diffusion of CaM-Ca^2+^ into condensates, suggesting selective diffusion rates of CaM in the absence and in the presence of Ca^2+^.

We integrated the above results using a cellular model of the CaM/Ca^2+^-dependent activation pathway of cMLCK (fig. S9). A detailed description of the model is given in the Methods section.

Using this model, we compared the effects of cMLCK localization (condensates vs cytosol) on the level of activation depending on the freely available CaM in the cardiomyocytes, assuming a total CaM concentration of 6 μmol L^-1^ (*28*) (Fig. 3J). Strikingly, only when cMLCK is localized to condensates can the kinase be significantly activated at physiological relevant Ca^2+^ and CaM concentrations (<10% of total pool).

### cMLCK condensates actively recruit co-factors and substrates

The results presented above strongly suggest that cMLCK is not released from condensates upon Ca^2+^/calmodulin activation, which leads to the question of how the kinase localized in condensates on the SR can phosphorylate its substrate myosin cRLC, which is primarily bound to the myofilaments. Using detergent permeabilization and fractionation of myocardial strips via differential centrifugation, we determined that about 5-10% of the total cRLC is soluble in myocardium from human organ donors (fig. S10A). Previous studies in animal models have shown that the cRLC rapidly exchanges between the myofilament and cytosolic space with a halftime of a few minutes (*29*). Overall, this suggests that a significant pool of soluble cRLC is available in the cytosol of cardiomyocytes for phosphorylation by cMLCK in the condensate phase near the SR, and that these phosphorylated cRLCs can subsequently be incorporated into the myofilaments.

Previous studies have shown that selective enrichment of client molecules is an emergent property of biomolecular condensates (*10, 12*). We hypothesized that cMLCK condensates selectively enrich substrates for the phosphorylation reaction and tested this hypothesis using co-condensation experiments with Alexa647 (A647) - labeled NTD and various potential client molecules labeled with Alexa546 (A546). cMLCK NTD condensates selectively enriched the isolated cRLC with a partition coefficient (PC) of about 5 (Fig. 4A). Identical results were obtained using the unlabeled NTD, suggesting that fluorophore-labeling did not affect client enrichment (fig. S10B). Similarly, co-expression of mTFP-labeled cRLC with cMLCK-FusionRed in COS-1 cells showed enrichment of cRLC in the condensates phase (fig. S10C). Next, to test if the condensates selectively enrich clients, we performed control experiments with both isolated bovine serum albumin (BSA) and cardiac troponin C (cTnC). The latter is highly homologous to cRLC with a high degree of structural similarity and biophysical properties. Surprisingly, BSA showed a PC below 1, suggesting that cMLCK NTD condensates actively reduce their internal concentration (Fig. 4B). cTnC showed a partition coefficient of close to 1, indicating very weak enrichment compared to cRLC. In contrast, cMLCK NTD condensates enriched Cy2-labeled ATP with a PC >2. These results show that cMLCK condensates can selectively enrich substrate molecules required for the phosphorylation reaction (i.e. cRLC and ATP).

**Figure 4.**
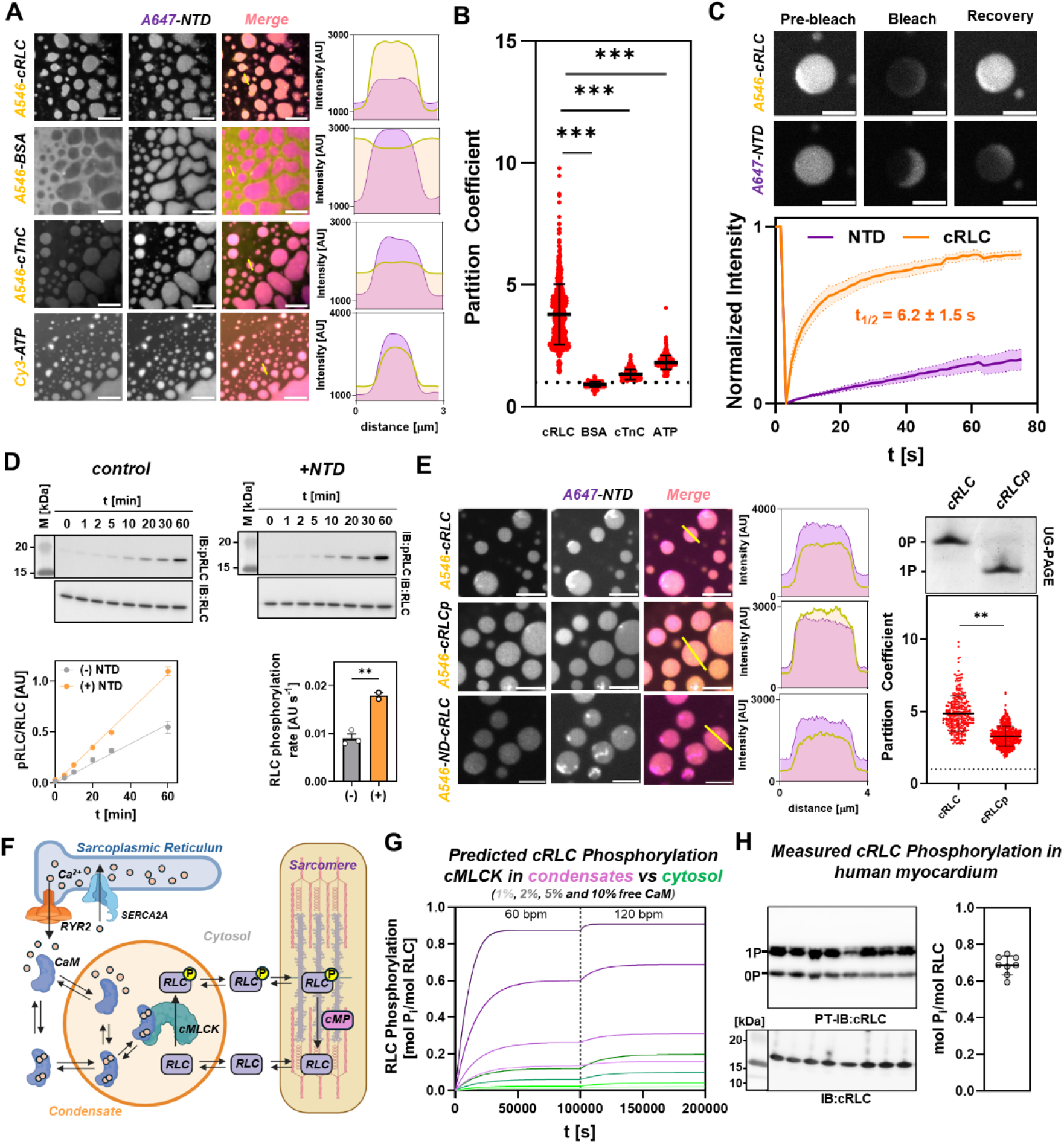
cMLCK condensates selectively enrich clients to increase catalytic efficiency. (A) Confocal micrographs of co-condensation of recombinant cMLCK NTD labeled with Alexa647 (A647) and various clients labeled with Alexa 546 (A546) or Cy2. Scale bar represents 5 μm. (B) Analysis of partition coefficients from (A) (pooled data from N=3-6). (C) FRAP analysis of cMLCK A657-NTD condensates in the presence of A546-cRLC (N=2, n=7). Scale bar represents 2 μm. (D) In-vitro kinase assay with recombinant cMLCK either in solution (top) or incorporated into biomolecular condensates (bottom) (N=2-3). (E) Confocal micrographs of co-condensation of cMLCK NTD with unphosphorylated (cRLC) and serine 15 phosphorylated cRLC (cRLCp), and N-terminally deleted cRLC (ND-cRLC). Phosphorylation levels were analyzed by urea-glycerol gel electrophoresis (UG-PAGE) (pooled data from N=3). (F) Schematic of the cellular model incorporating cRLC exchange between sarcomeres, cytosol and condensates (RYR2 – ryanodine receptors, SERCA2a-sarcoplasmic reticulum calcium ATPase 2a; CaM-calmodulin; RLC-myosin regulatory light chain; cMP–cardiac myosin phosphatase). (G) Model predictions of the level of cRLC phosphorylation depending on available calmodulin and heart rate for cMLCK in condensates or freely diffusible in the cytosolic phase. (H) cRLC phosphorylation levels in myocardium from human donors (N=8).

Efficient phosphorylation requires rapid exchange of products with new substrate molecules in the close vicinity of the protein kinase. We measured the dynamics of cRLC exchange between the buffer and condensates phase using FRAP, which showed a halftime of recovery of the cRLC signal in the condensates of about 6 s (Fig. 4C). This demonstrates that although the condensates can highly enrich cRLC, they also rapidly exchange it with the local environment and therefore facilitate the phosphorylation reaction. Indeed, embedding the cMLCK in condensates significantly increased its catalytic activity towards isolated cRLC (Fig. 4D). Moreover, we tested if the product of the kinase reaction is similarly enriched in condensates phase by co-condensation of the cMLCK NTD with serine 15 phosphorylated cRLC (Fig. 4E). Strikingly, the phosphorylated form of the cRLC showed a significantly lower PC compared to the unphosphorylated cRLC, suggesting that electrostatic interactions of the N-terminal extension of cRLC regulate its interaction with the condensates. In excellent agreement, removal of the unique N-terminal extenstion of the cRLC (ND-cRLC) significantly reduced its partition coefficient into the condensate.

We integrated these data into the cellular model described above to determine whether cMLCK in condensates can phosphorylate cRLC to significant levels (Fig. 3J and Fig. 4F). We determined the level of cRLC phosphorylation in the myofilaments as a function of heart rate and available calmodulin concentration (Fig. 4G) and compared the values to those of cRLC phosphorylation measured in human myocardium from organ donors (Fig. 4H). The model predicts that even low levels of available calmodulin (1-5%) lead to significant levels of cRLC phosphorylation (about 0.7 mol P_i_ mol RLC^-1^) approaching the values measured in human myocardium if cMLCK is organized into biomolecular condensates (Fig. 4H). Strikingly, the model further predicts that less than 1% of the total cRLC pool is required to be outside the sarcomere at any given time point, significantly less than the level measured from human organ donors (fig. S10A). In contrast, if cMLCK is allowed to freely diffuse in the cytosol and sarcomere space, the level of cRLC phosphorylation never exceeds 0.2 mol P_i_ mol cRLC^-1^ in the simulations. These results are consistent with a model in which myosin motors are not phosphorylated in-situ by cMLCK but rather a small pool of soluble cRLC is rapidly phosphorylated by cMLCK in condensates and incorporated into the myofilaments.

## Discussion

Biomolecular condensation of protein kinases is emerging as a new paradigm to understand cellular signaling in both health and disease states (*10, 30*). Condensation compartmentalizes kinases, their substrates and required cofactors, modulates substrate selectivity and allows spatiotemporal control of signaling events (*31*). Phase separation is already an established fundamental concept for kinase function in cancer cell biology and neurological disorders (*32*) but is understudied in the context of heart muscle contractile regulation. The present study extends this framework to cardiac muscle physiology, thus establishing a previously unrecognized mechanism for the spatiotemporal regulation of myosin motor activity in the heart.

Cardiac myosin light chain kinase spontaneously undergoes LLPS controlled by its unique intrinsically-disordered N-terminal extension (Fig. 2). cMLCK condensates are associated with the intracellular Ca^2+^ storage compartments of human cardiac muscle cells (Fig. 1), allowing precise control of its kinase activity via the Ca^2+^-dependent recruitment of calmodulin (Fig. 3). Ca^2+^/calmodulin-activated cMLCK remains in the condensate space, which can selectively recruit and enrich substrate molecules (i.e. ATP and cRLC), thus increasing its catalytic efficacy (Fig. 4D). Our simulations clearly demonstrate that the low intrinsic activity of cMLCK combined with the low availability of free calmodulin in cardiomyocytes is in inconsistent with cMLCK operating as a soluble kinase (Fig. 4G). These results lead to a novel concept of cardiac myosin motor regulation by cMLCK (Fig. 4F). The majority of cRLCs are bound to the myosin motors in the thick filaments but in a rapid equilibrium with a small pool of free soluble cRLCs in the cytosol, which might be newly synthesized or released from the myofilaments (fig. S10A). The free cRLC is recruited into SR-associated cMLCK condensates, where it is phosphorylated in a Ca^2+^-dependent manner and released back into the cytosol. The cytosolic phosphorylated cRLCs are subsequently exchanged into the myofilaments and subject to dephosphorylation via myofilament-bound myosin phosphatase enzyme complexes.

This type of regulation is fundamentally different than that observed in both skeletal and smooth muscle, likely reflecting the different functional requirements of these types of muscles. Smooth muscle contraction is directly activated by myosin motor phosphorylation (*33*). In contrast, RLC phosphorylation has been suggested as the molecular basis of post-tetanic potentiation of skeletal muscle contractile function (*34*). Both processes involve rapid changes in the level of myosin motor phosphorylation associated with cellular calcium activation and operate in a ‘switch-like’ fashion (*9*). This is facilitated by the high affinity for calmodulin and catalytic activity of both skeletal and smooth muscle MLCKs. In contrast, the low intrinsic activity of MLCK is unique to cardiomyocytes and (*35*), in combination with biomolecular condensation, likely enables spatiotemporal fine control over the level of motor phosphorylation and cardiac output on a timescale longer than the individual Ca^2+^ transient.

Both altered myosin motor activity and reduced availability have been suggested as the molecular etiology of genetic cardiomyopathies (*36*), and lower levels of myosin motor phosphorylation have been associated with impaired contractile function during heart failure (*6, 37*). It has previously been shown that dysregulation of the physicochemical properties and dynamics of biomolecular condensates drives an expanding class of human pathologies, colloquially called ‘condensatopathies’ (*30, 38*). We hypothesize that dysregulated biomolecular condensation of cMLCK might either present as accummulation of hardened cardiotoxic aggregates or alternatively as failed LLPS, leading to low cMLCK activity and lower levels of myosin phosphorylation in heart disease.

As with any experimental system, we acknowledge several limitations in the current study. Overexpression of labeled proteins and fusion with fluorescent tags might have affected the LLPS behavior of cMLCK, although unperturbed phase separation of the untagged cMLCK NTD suggests little to no effect of fluorophore attachment (fig. S5B). In-vitro or in-cellula phase separation of isolated proteins allows precise control of experimental conditions such as stoichiometries, buffer conditions, etc., but does not fully recapitulate the in-vivo condition. Mechanical forces and additional condensate components such as lipids, RNA or DNA may alter condensation behavior inside cells.

Viewed more broadly, our study has uncovered an underappreciated mechanism for myofilament contractile regulation via the control of protein kinase function by biomolecular condensation. Physiological and patho-physiological relevant protein kinases in the heart such as protein kinase A (PKA) and calmodulin-dependent protein kinase II (CamKII) have been shown to be regulated via LLPS in different cell systems, suggesting similar mechanisms might operate in the heart (*39, 40*). The identification of cMLCK condensate formation as a regulatory node in cardiac myosin phosphorylation suggests that the biophysical properties of the condensate microenvironment represent a potential therapeutic target for restoring normal myosin phosphorylation levels in heart failure (*41*).

## Materials and Methods

Detailed description of the materials and methods can be found in the supplementary information.

## Acknowledgments

The authors thank the Light Microscopy Core at the University of Kentucky for instrumentation access and technical support (RRID:SCR_026405).

## Funding

National Institutes of Health grants R01HL173989 (to TK, KSC); R01HL148785 (to TK, KSC); American Heart Association grant 25PBA1503383 (to TK)

## Author contributions

Conceptualization: TK; Methodology: TK, KJC, EE, XF; Investigation: KJC, AWH, IRS, EE, TK; New reagents: KSC; Visualization: XF, TK; Funding acquisition: KSC, EE, TK; Project administration: TK; Supervision: TK; Writing – original draft: KJC, AWH, IS, EE, TK; Writing – review & editing: KJC, AWH, IS, EE, TK.

## Competing interests

Authors declare that they have no competing interests.

## Data, code, and materials availability

All data are available in the main text or the supplementary materials. Human tissue requires an MTA agreement. All materials are available at reasonable request from the corresponding author.

## Supplementary Materials

Supplementary Text

Materials and Methods

Figs. S1 to S10

Tables S1 to S2

Supplementary References

## Notes

### Competing Interest Statement

The authors have declared no competing interest.

