## Supplementary Information for "Biomolecular condensation of cMLCK enables myosin motor phosphorylation in the heart"

### Supplementary Text

#### Estimation of cMLCK concentration and cluster size in ventricular myocardium

100 mg human ventricular tissue was homogenized in 1 mL of extraction buffer (please see Methods for details). The density of human myocardium was assumed to be 1.055 g $mL^{-1}$, giving a total of 95 µL of myocardium per mL extraction buffer. The fraction of cardiomyocytes in the mid-ventricular wall is about 70% (v/v) and about 10% (v/v) of the cardiomyocyte volume is occupied by the cytosol, resulting in 6.6 µL of cardiomyocyte cytosol dissolved in 1 mL extraction buffer. 8 µL of extraction buffer were loaded onto SDS-PAGE per donor sample, which is equivalent to 0.053 µL of cardiomyocyte cytosol volume. The average amount of cMLCK in the samples was estimated by linear regression of the protein standards to 0.189 pmol (Fig. S1B), giving a cMLCK concentration of 0.189 pmol 0.053 $\mu L^{-1}$ cytosol, which is equivalent to 3.6 µmol $L^{-1}$.

The average sarcoplasmic reticulum surface area to cardiomyocyte volume is about 1.19 $\mu m^2$ $\mu m^{-3}$ (*1*). The average number of cMLCK clusters per $\mu m^2$ of sarcoplasmic reticulum can be calculated to 1.56 cluster $\mu m^{-2}$ from the average nearest neighbor distance assuming a Poisson distribution, giving a total of 1.86 cMLCK clusters per $\mu m^3$ cardiomyocyte volume. The concentration of cMLCK in cardiomyocytes was estimated to about 3.6 µmol $L^{-1}$ (Fig. S1B), which is equivalent to 2168 molecules per $\mu m^3$, giving an average of 1065 cMLCK molecules per cluster.

### Materials and Methods

#### Human myocardial samples and protocols

All procedures were approved by the University of Kentucky Institutional Review Board (IRB#46103), and the subjects or their legally authorized representative gave written informed consent. A detailed protocol for the procurement collection, processing and storage of human myocardial tissue was published previously (*2*).

#### Tissue processing

*Cryosectioning:*
Cardiac tissue specimens were obtained from a biobank and maintained on dry ice throughout processing to preserve structural integrity. Samples were removed from cryogenic vials, trimmed using a sterile razor blade and forceps to generate appropriately sized fragments. A small volume of optimal cutting temperature (OCT) compound was applied to anchor the tissue, followed by complete embedding to ensure uniform support. Cryosectioning was performed on a cryostat using a microtome, with blade integrity verified to minimize sectioning artifacts. Sections (7 μm thickness) were affixed to SuperFrost Plus (VWR) microscope slides by gentle adhesion, avoiding direct compression. Slides were kept at -20 °C until immunostaining was performed.

*Myofilament fractionation:*
50 mg frozen mid-ventricular samples were homogenized in myofibril buffer (20 mmol $L^{-1}$ MOPS pH 7, 100 mmol $L^{-1}$ NaCl, 2 mmol $L^{-1}$ $MgCl_2$, 1 mol $L^{-1}$ EGTA, 1:20 dilution Triton X-100 with added PhosStop (Roche) phosphatase inhibitor cocktail and cOmplete Mini (Roche) EDTA-free protease inhibitor cocktail). The samples were centrifuged for 5 minutes at 17,500 rpm, 4°C. The supernatants were transferred to pre-chilled microtubes (soluble fraction). The pellets were subsequently rewashed with myofibril buffer and centrifuged under the same conditions. The supernatants were discarded, and the volumes of the remaining insoluble fractions were adjusted with 10mmol $L^{-1}$ Tris to match those of the soluble fractions. 5x SDS-PAGE loading buffer (250mmol $L^{-1}$ Tris, 10% (w/v) SDS, 30% (v/v) glycerol, 0.05% (w/v) bromophenol blue, and 500mmol $L^{-1}$ DTT) was added to achieve a final 1x concentration. The samples were denatured at 100°C for 2 minutes, then rapidly frozen in liquid nitrogen and stored at –80°C.

*Sarcoplasmic reticulum fractionation:*
100 mg frozen mid-ventricular samples from non-failing organ donors were homogenized three times for 10 seconds in 500 μL homogenization buffer (10 mmol $L^{-1}$ Tris pH 8, with added cOmplete Mini EDTA-free protease inhibitor cocktail (Roche)). Cell debris was removed through two centrifugation steps, each lasting 15 minutes at 17,500 rpm and 4°C. The supernatants were transferred to fresh, pre-chilled microtubes following each procedure. The supernatants were then sedimented for 30 minutes at 30,000g. The resulting supernatants, representing the cytosolic fraction, were carefully transferred to fresh microtubes and diluted with 120 μL of 5x SDS-PAGE running buffer. The pellets were resuspended in resuspension buffer (600 mmol $L^{-1}$ KCl pH 7, and 30 mmol $L^{-1}$ histidine) followed by sedimentation as previously described. The supernatants were discarded and the pellets, representing the sarcoplasmic reticulum fraction, were resuspended in 500 μL of

1x SDS-PAGE running buffer. Both the cytosolic and sarcoplasmic reticulum fractions were denatured at 100 °C for 2 minutes, then rapidly frozen in liquid nitrogen and stored at –80 °C.

**Western-blot analysis**

SDS-PAGE samples were analyzed using 4–20% (v/v) acrylamide gradient gels (Invitrogen) and subsequently transferred to nitrocellulose membranes using the iBlot3 rapid-dry transfer system (ThermoFisher). Membranes were blocked in either 5% (w/v) non-fat dried milk or 5% (w/) bovine serum albumin in TBS-T (50 mmol $L^{-1}$ Tris, 150 mmol $L^{-1}$ NaCl, 0.1% Tween-20, pH 7.4) for 1 h at room temperature. Primary antibodies were incubated overnight at 4°C. HRP-conjugated secondary antibodies were applied for 1 h at room temperature and detected by enhanced chemiluminescence ECL (Pierce). A full list of antibodies (including dilutions) is given in Table S1. Images were acquired on a ChemiDoc imaging system (Bio-Rad). For Phos-tag immunoblotting, gels were prepared with 50 µmol $L^{-1}$ Phos-tag acrylamide and 100 µmol $L^{-1}$ $MnCl_2$ according to the manufacturer's instructions (Wako Chemicals).

**Plasmid constructs**

All plasmids were purchased from GenScript and all sequences were verified by sanger sequencing (Tab. S1). Mammalian expression constructs were generated by inserting the optimized sequences for expression in COS-1 cells into a pcDNA3.1(+) vector using the BamHI and EcoRV site. For bacterial expression of constructs, codon optimized sequences were inserted into either the pET3a or pET11a vector using the NdeI/BamHI or NheI/BamHI restriction sites, respectively. Plasmid constructs are summarized in Table S2.

**Cell culture**

COS-1 cells (Sigma) were cultured in maintenance medium (10% (w/v) fetal calf serum, 1% (w/v) penicillin–streptomycin in high-glucose DMEM) at 37°C in 5% (v/v) $CO_2$ in a HeraCell i150 incubator. For transfection, cells were plated onto 35-mm dishes and grown to 50–70% confluency. COS-1 cells were transfected with 1 µg of plasmid DNA using ESCORT IV (Sigma) transfection reagent in maintenance medium without fetal calf serum according to manufacturer instructions and cells were transferred into maintenance medium the next morning. For live cell imaging, COS-1 cells were plated in 35 mm glass bottom dishes and transfected as described above. Prior to live cell imaging, maintenance medium was replaced with FluoroBrite DMEM (ThermoFisher) containing 1% (w/v) penicillin–streptomycin. Cells were maintained at 37°C and 5% (v/v) $CO_2$ throughout the imaging using a live-cell imaging chamber.

Primary cultures of neonatal rat cardiomyocytes were generated as described previously by collagenase digestion of newborn rat hearts (*3*). Isolated cells were plated onto collagen-coated 35 mm plastic dishes (Falcon Primaria) in Plating Medium (33% (v/v) DMEM, 16% (v/v) M199, 10% (v/v) horse serum, 5% (v/v) fetal calf serum, 4 mmol $L^{-1}$ glutamine, 1% (v/v) penicillin-streptomycin) over-night and switched to transfection medium (73% (v/v) DBSS-K, 21% (v/v) M199, 4% (v/v) horse serum, 4 mmol $L^{-1}$ glutamine; DBSS-K: 116 mmol $L^{-1}$ NaCl, 1 mmol $L^{-1}$ $NaH_2PO_4$, 0.8 mmol $L^{-1}$ $MgSO_4$, 5.5 mmol $L^{-1}$ glucose, 32.1 mmol $L^{-1}$ $NaHCO_3$, 1.8 mmol $L^{-1}$ $CaCl_2x2H_2O$, pH 7.2) the next morning. Transient transfections with plasmid DNA were carried out over-night using Escort III (Sigma) as transfection agent and the cells were transferred into Maintenance Medium (73% DMEM, 20% M199, 4% horse serum, 4 mmol $L^{-1}$ glutamine, 1% penicillin-streptomycin, 0.1 mmol $L^{-1}$ phenylephrine, 10 μmol $L^{-1}$ cytosine arabinoside) the next morning.

**Immunofluorescence**

*Immunostaining of cryosections*

The slides with adhered tissue samples were removed from –20°C storage and allowed to come up to room temperature. The tissue samples were encircled with an ImmEdge (Vector Labs) hydrophobic barrier pen. A solution of 4% (v/v) paraformaldehyde (PFA) in 1x phosphate buffered saline (PBS) was added to completely cover the samples to incubate for 5 minutes. The PFA solution was aspirated, and the samples were washed three times with PBS, each sitting for 5 minutes. A solution of 0.5% (v/v) Triton X-100 in PBS was added to cover the samples for 5 minutes. The Triton solution was aspirated, and the samples were washed three times with PBS, each sitting for 5 minutes. For staining of sarcoplasmic reticulum proteins, cryosections were dried at room temperature for 12 h, fixed for 5 mins in acetone at -20°C, washed three times in PBS and permabilized for 5 mins using 0.05 % (wv) saponon (Sigma) in PBS. The primary antibody solution of 1x Gold Buffer and specific antibodies (see Table S2 for details) was prepared, added to cover the samples, and incubated in a chamber with constant humidity at 4°C overnight. Following this, the primary antibody solution was aspirated, and the samples were washed three times with PBS, each sitting for 5 minutes. The secondary antibody solution of 1x Gold Buffer and specific antibodies (see Table S1 for details) was prepared, added to cover the samples, and incubated as before at room temperature on a rocker for 1 hour. Following this, the secondary antibody solution was aspirated, and the samples were washed three times with PBS, each sitting for 5 minutes. Following the final wash, Lisbeth media (1 mol $L^{-1}$ Tris, 90% (v/v) glycerol, 235 mmol $L^{-1}$ n-propyl-gallate) was added dropwise onto the samples. A coverslip was added onto the samples, where its edges were sealed to the slide with TissueTek VIP (Sakura) wax. The slides were stored at 4°C. Images were taken on a Nikon CSU-W1 SoRa Confocal Microscope with super-resolution capabilities using a 60x/1.4 NA oil immersion objective.

*Immunostaining of COS-1 cells*

Cells were washed three times with PBS and fixed with 4% paraformaldehyde in PBS for 5 minutes at room temperature. Following fixation, cells were washed three times in PBS (5 minutes each) and permeabilized with 0.1% Triton X-100 in PBS for 3–5 minutes. Cells were then washed three times in PBS before incubation with primary antibodies diluted 1:100 in 1× Gold buffer. Mouse anti-p62/SQSTM1 and rabbit anti-MYLK3 antibodies were applied either individually or in combination and incubated overnight at 4 °C in a humidified chamber. After primary incubation, cells were washed three times in PBS and incubated with fluorophore-conjugated secondary antibodies (goat anti-mouse Alexa Fluor 594 and goat anti-rabbit Alexa Fluor 488, each at 1:100 in 1× Gold buffer) for 1 hour at room temperature in the dark. Cells were subsequently washed three times in PBS and mounted using an anti-fade mounting medium. Samples were allowed to equilibrate to room temperature prior to imaging. Images were taken on a Nikon CSU-W1 SoRa Confocal Microscope with super-resolution capabilities using a 60x/1.4 NA oil immersion objective.

*Immunostaining of neonatal rat cardiomyocytes*
After 48 hours, transiently transfected NRC cultures were washed once with PBS, fixed for 10 min at RT in 4% (v/v) PFA/PBS for 10 minutes and subsequently permeabilized with 0.2% (v/v) Triton X-100/PBS for 5 min. Primary and secondary antibodies were diluted in 1%BSA/GB as described previously (*4*). Images were taken on a Leica SP5 Confocal Microscope, equipped with a blue diode, argon and helium-neon lasers using a 63x/1.4 NA oil immersion lens.

**Fluorescence life-time and fluorescence recovery after photobleaching imaging**

Fluorescence life time imaging was performed on a PicoQuant FLIM system mounted on a Nikon A1 confocal microscope with 60x water-immersion objective. Time-correlated single photon counting (TCSPC) data were acquired using a PicoHarp 300 module and a tunable white light laser (NKT Photonics, SuperK) operated at a repetition rate of 18.7 MHz with 488nm excitation. Fluorescence lifetime data were collected and analyzed using SymPhoTime 64 software (PicoQuant).

Fluorescence recovery after photobleaching (FRAP) experiments were performed on a Nikon AXR Inverted Confocal Microscope equipped with a 60× oil-immersion objective. A circular region of interest (ROI) with a radius of 1 µm was photobleached using a 405 nm laser at 15–40% laser power for 10 iterations. Pre-bleach images were acquired prior to bleaching, and fluorescence recovery was monitored using a 488 nm laser at a frame rate of 1 frame $s^{-1}$. Time-lapse images were collected for 300 seconds post-bleach. Fluorescence recovery within condensates was quantified using Fiji (ImageJ 1.54p), and normalized recovery curves were fitted using GraphPad Prism 10.

**Image analysis**

Images were exported as nd2 files and loaded into Fiji (ImageJ 1.54p) (*5*). Condensate sizes and shape were analyzed using the built-in particle analysis option in ImageJ after binarizing the image using appropriate threshold settings and watershed separation. Partition coefficients were determined by measuring the mean intensity value of the condensates and adjacent dilute phase. Colocalization was determined using the Coloc2 plugin. Cluster and nearest neighbor analysis were performed using the spatial analysis option of the 3D ImageJ Suite and the Mosaic Interaction Analysis plugin (*6, 7*).

**Protein preparation**

Proteins were expressed in chemically competent BL21(DE3) RIPL cells (Agilent Technologies) using the bacterial expression plasmids summarized in Table S1. Protein expression was induced with 0.2-1 mmol $L^{-1}$ isopropyl-β-D-thiogalactoside (IPTG) once bacterial cultures reached an $OD_{600\ nm}$ of about 0.6 and incubated overnight at either 18°C (for NTD) or 30°C (for RLC and cTnC), depending on the protein construct. The next morning cells were harvested by centrifuged for 8 minutes at 9,500rpm at 4°C. The pellets were washed with phosphate buffered saline (PBS) and centrifuged again as before. The supernatants were discarded and pellets were snap frozen in liquid nitrogen and stored at –80°C until further use.

*Purifcation of cMLCK variants*

HisTrapFF (Cytiva) columns were equilibrated with 15 column volumes (CV) wash buffer (25 mmol $L^{-1}$ HEPES, 300 mmol $L^{-1}$ NaCl, 10 mmol $L^{-1}$ imidazole, 6 mol $L^{-1}$ urea, pH 7.5, sterile filtered using a 0.45 µm syringe filter), then with 15 CV elution buffer (wash buffer containing 500 mmol $L^{-1}$ imidazole), and then with 15 CV wash buffer again. The bacterial pellets were thawed at room temperature, then lysed with 20 mL BugBuster Protein Extraction Reagent (MilliporeSigma) containing 5 mg $mL^{-1}$ bovine deoxyribonuclease I (Thermo Scientific) and 10 mg $mL^{-1}$ chicken lysozyme (MP Biomedicals) on ice and rocked for 10 minutes. Subsequently 5 mL of the wash buffer was added and the lysate was centrifuged for 8 minutes at 9,500 rpm, 4°C. The supernatant was sterile filtered through a 0.45 µm filter, flowed over the columns, and then collected as the flowthrough fraction. The columns were washed with 15 CV of wash buffer and proteins eluted with 15 CV of elution buffer. The elution fractions were combined, 5% (v/v) glycerol was added to stabilize the protein, and the sample was centrifuged for 8 minutes at 9,500rpm, 4°C. The sample was concentrated to roughly 4 mL using a 10 kDa cutoff spin concentrator (Sartorius) at 5000 g and 4°C. The sample was dialyzed overnight in dialysis buffer (20 mmol $L^{-1}$ HEPES, 300 mmol $L^{-1}$ NaCl, 5% (v/v) glycerol, 1 mmol $L^{-1}$ dithiothreitol, pH 7), using 1 mL 3.5-5kDa Float-A-Lyzer G2 (Spectra-Por) dialysis devices. The sample concentration was determined via UV absorbance using calculated extinction coefficients at 280 nm and a NanoDrop One spectrophotometer (Thermo Fisher). Protein aliquots were snap frozen in liquid nitrogen and stored at –80°C until further use.

*Purification of cRLC/cTnC variants*

The human ventricular myosin regulatory light chain (cRLC) and cardiac troponin C (cTnC) were purified as described previously (*8, 9*). Proteins were labeled with Alexa647-NHS or Alexa564-NHS (ThermoFisher) according to manufacturer's instructions. Briefly, proteins were buffer exchanged into 200 mmol $L^{-1}$ sodium bicarbonate buffer pH 8.5 using NAP5 columns and fluorophores were added from a DMSO stock solution to 1:1 stoichiometry. Reactions were incubated at room temperature in the dark for 60 mins, quenched by adding an excess of glycine pH 7 and free dye removed by gel-filtration using NAP10 columns. Proteins were concentrated using 10 kDa cutoff spin concentrators, aliquoted, snap frozen in liquid nitrogen and stored at -80°C until further use.

**In-vitro kinase assay**

Recombinant human cRLC was gel-filtered into cMLCK assay buffer (composition in mmol $L^{-1}$: 50 HEPES, 50 KCl, 1 $MgCl_2$, 1 $CaCl_2$, 1 DTT) using NAP5 columns (GE Healthcare) and protein concentration determined by UV absorbance using calculated extinction coefficients at 280 nm. Concentration of cRLC was adjusted to 20 µmol $L^{-1}$, and recombinant calmodulin and cMLCK were added to final concentration of 0.5 µmol $L^{-1}$ and 0.1 µmol $L^{-1}$, respectively. For some assays recombinant isolated NTD was added to a final concentration of 50 µmol $L^{-1}$ to test for the effects of phase separation. cRLC phosphorylation was analyzed by Western-blot using a phospho-specific and total antibody (Tab. S2), or urea glycerol PAGE as described previously (*10*).

**PIP strip assay**

PIP strips (Echelon Biosciences) were blocked with 1% non-fat milk powder in PBS for 1 hr followed by incubation with 2 µg $mL^{-1}$ NTD for 1 hr in PBS containing 1% non-fat milk powder at room temperature. Subsequently, PIP strips were washed three-times for 5 mins in PBS containing 0.05% (v/v) Tween-20 (PBS-T) and incubated with primary and secondary antibody as described above. PIP strips were developed with enhanced chemiluminescence ECL (Pierce).

**Protein interaction analysis**

Microscale thermophoresis binding experiments were performed on Monolith X (Nanotemper, Germany) using premium capillaries. Proteins were gel-filtered into assay buffer (composition in mmol $L^{-1}$: 10 Tris-HCl, 100 NaCl, 2 DTT) prior to experiments. Alexa 647-labeled proteins were used at a fixed concentration of 50 nmol $L^{-1}$.

Analytical gel filtration experiments were performed on Superdex 75 100/300 HR column (Cytiva) in PBS containing 1 mmol $L^{-1}$ DTT at a constant flow rate of 0.5 mL $min^{-1}$.

**Phase separation assays**

NTD was diluted from a stock solution into assay buffer (10 mmol $L^{-1}$ Tris HCl pH 7, 50 mmol $L^{-1}$ NaCl, 1 mmol $L^{-1}$ DTT) to a final concentration of 50-100 µmol $L^{-1}$ and mixed with Alexa 647-labeled NTD to a final stoichiometry of 20:1. A small droplet (20 µl) was added to a microscope coverslip (VWR, #1) and condensate formation was monitored via confocal microscopy using both Alexa 647 fluorescence and phase contrast. For co-condensation experiments, 1 µmol $L^{-1}$ of Alexa 546 labeled proteins of Cy2-labeled ATP were added and monitored using confocal microscopy with appropriate excitation and emission filter settings.
Coverslips were silanized by cleaning them with diluted 2% (v/v) Hellmanex III solution (VWR Avantor) for 20 mins followed by extensive washing in MilliQ water to remove the cleaning solution and drying under nitrogen stream. Sigmacote solution (Sigma) was directly added to cleaned coverslips, incubated for 2 mins at room temperature and removed by aspiration. Coated coverslips were washed with MilliQ water and dried a fume hood.

**Preparation of giant unilamellar vesicles**

Giant unilamellar vesicles were prepared as described previously using the water-in-oil elmusion transfer method (*11*). Briefly, both 1,2-dioleoyl-sn-glycero-3-phosphocholine (DOPC, Sigma) and ATTO 488-labeled 1,2-dioleoyl-sn-glycero-3-phosphoethanolamine (ATTO488-DOPE, Sigma) were dissolved in a 2:1 mixture of chloroform and methanol to 10 mg $mL^{-1}$. Lipids were mixed to a final concentration of 0.1% (mol $mol^{-1}$) ATTO488-DOPE and diluted into 400 µL mineral oil to a final concentration of 0.1 mg $mL^{-1}$. The solution was heated to 80°C in an open microcentrifuge tube for 15 mins to completely dissolve the lipids and remove any residual chloroform and methanol. 40 µL 10 mM Tris-HCl pH 7, 100 mmol $L^{-1}$ NaCl, 100 mmol $L^{-1}$ sucrose, 1 mmol $L^{-1}$ DTT were added to the mineral oil solution and the mixture vigorously vortexed for 40 s and stored at 4ºC for 15 mins to equilibrate. 100 µL 10 mM Tris-HCl pH 7, 100 mmol $L^{-1}$ NaCl, 100 mmol $L^{-1}$ glucose was added to a fresh microcentrifuge tube and carefully overlayed with the mineral oil/buffer suspension, incubated for 10 mins at room temperature and centrifuged at 16,000 g for 10 mins to harvest the GUVs. After removal of the mineral oil and most of the aqueous layer, the

GUV pellet was resuspended in fresh buffer and centrifuged again at 16,000 g for 10 mins. The resulting pellet was resuspended in 100 mL fresh buffer and immedeat;ey used for experiments.

**Biochemical modelling**

Biochemical models of cMLCK were created using COPASI v4.46 (Build 300) (*12*). The calcium transient was modelled as described previously using a two compartment system (*13*). The total concentration of calmodulin in cardiomyocytes was assumed to be 6 µmol $L^{-1}$ (*14*). The concentration of RLC in the sarcomere was estimated to 200 µmol $L^{-1}$, assuming sarcomeres occupy 60% of the cardiomyocyte space (*15*). Kinetic constant for $Ca^{2+}$ binding and unbinding from calmodulin were taken from (*16*). The kinetic constants for RLC dissection from thick filaments was taken from (*17*) with a steady state dissociation constant of 3 µmol $L^{-1}$ (*18*). The Michaelis-Menten constants for cRLC phosphorylation by cMLCK were taken from (*19*). Dephosphorylation kinetics of cRLC in intact hearts was taken from (*20*). All other parameters were experimentally determined in this study.

**Statistics**

Data are represented as mean ± standard deviation (s.d.) for parametric and median ± interquartile range for lognormal distributed data sets, with N and n referring to the number of biological and technical repeats, respectively. A detailed description of the statistical analysis of data is given in the associate figure captions with the following considered statistically significant: *p<0.05, **p<0.01, ***p<0.001, ****p<0.0001.

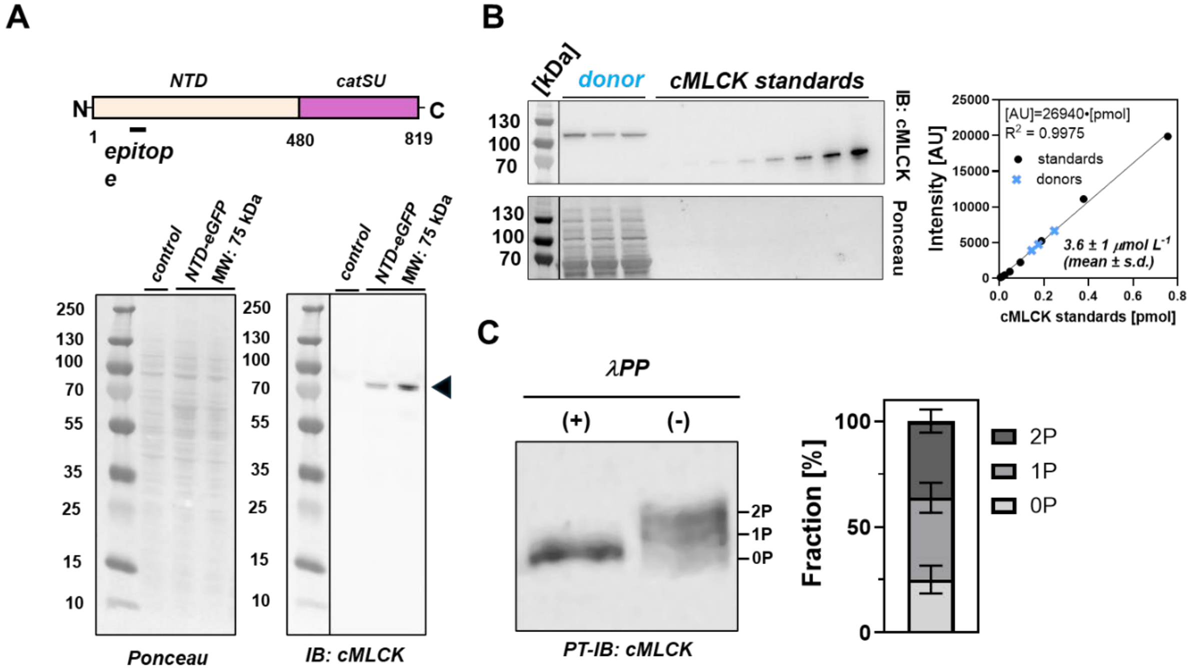


**Fig. S1. Validation of cMLCK antibody, estimation of cMLCK content and phosphorylation level.** (A) Top: Domain organization of cMLCK with antibody epitope indicated accordingly. Bottom: Immuno-blot of untransfected COS-1 cells and COS-1 cells transfected with plasmid expressing NTD-eGFP construct (estimated molecular weight is about 75 kDa). (B) Quantitative immune-blotting to estimate the cMLCK concentration in human ventricular cells. cMLCK standards used recombinantly expressed N-terminal domain (NTD) of cMLCK. (C) Phostag-immuno-blot (PT-IB) analysis of cMLCK phosphorylation levels in human donor myocardium. As a control (+), myocardial lysates were either treated with lambda protein phosphatase (λPP) to dephosphorylate cMLCK and identify the 0P band. Mean ± s.d. from N=4 human heart donor samples.

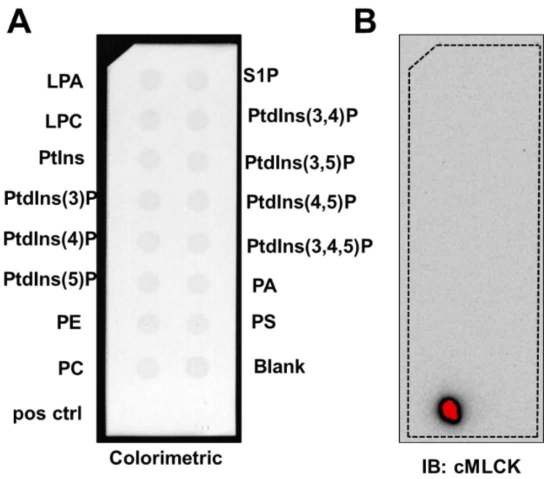


**Fig. S2. PIP strip assay with isolated recombinant cMLCK NTD.** (A) Colorimetric scan of the PIP strip. (B) Immuno-blot after incubation with recombinant cMLCK NTD. Please note that only the positive control gave a reproducible signal.

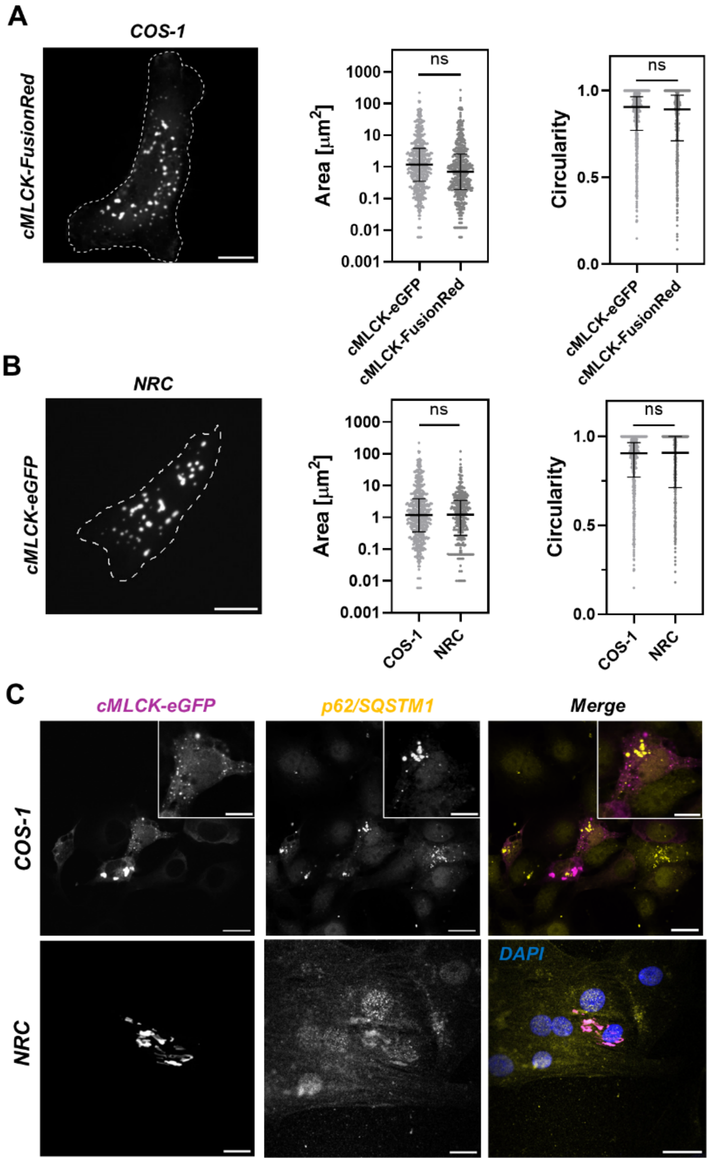


**Fig. S3. Effect of fluorescent-tag and cell type on cMLCK condensate formation. (A)** Left: Live cell image of COS-1 cells expressing FusionRed-labeled cMLCK. Right: Size and shape distribution of eGFP- and FusionRed-labeled cMLCK condensates. Data were analyzed with a nested two-tailed unpaired student's t-test (pooled data from N=3-5, n=20-50). (B) Left: Live cell image of NRC expressing cMLCK-eGFP. Right: Size and shape distribution of cMLCK-eGFP condens ates in COS-1 and NRCs. Data were analyzed with a nested two-tailed unpaired student's t-test (pooled data from N=2-5, n=20-50 cells). (C) Co-localization analysis of cMLCK-eGFP and p62/SQSTM1 in COS-1 cells and NRCs.

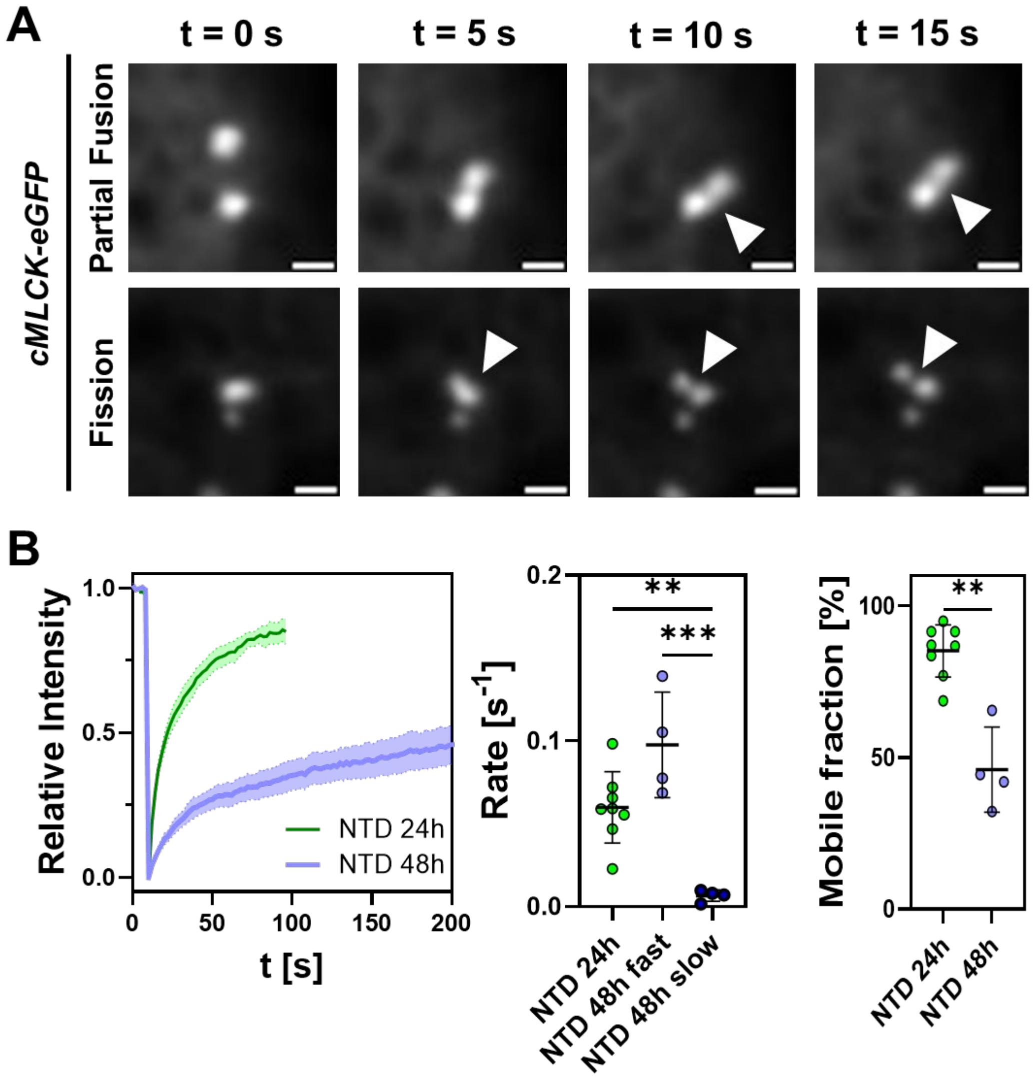


**Fig. S4. Dynamics of cMLCK-eGFP and NTD-eGFP condensates in COS-1 cells.** (A) Confocal images of partial fusion and fission events of cMLKC-eGFP condensates. (B) FRAP analysis of NTD-eGFP condensates 24h and 48h after COS-1 cell transfection. Data are mean ± s.d. for N=2-3 and n=4-8).

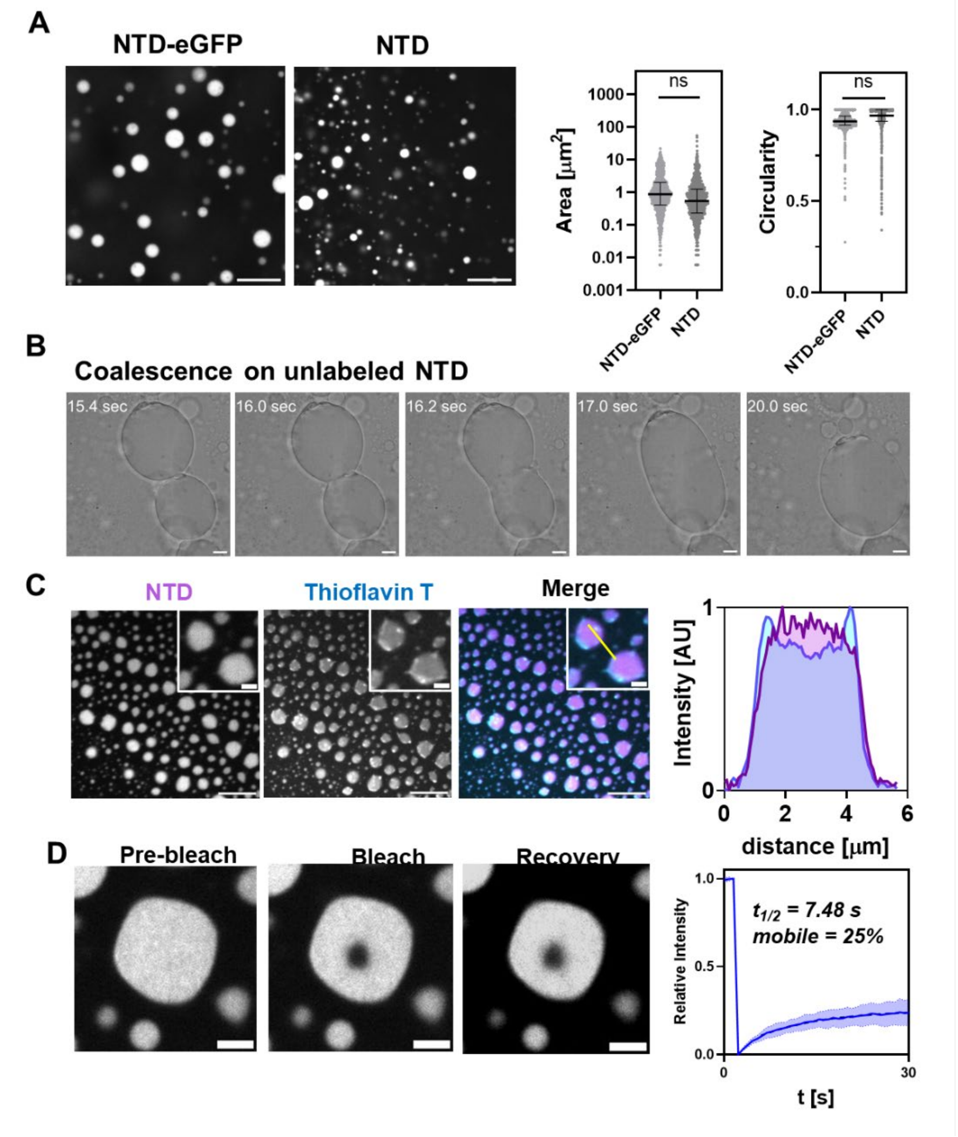


**Fig. S5. Characterization of isolated recombinant NTD condensates.** (A) Left: Confocal micrographs of condensates formed by NTD fused to eGFP (NTD-eGFP) or NTD mixed with 5% (mol mol$^{-1}$) Alexa 647 labeled NTD. Size and shape comparisons are shown on the right. Pooled data (median ± interquartile range) for N=3-6. (B) Coalescence of condensates formed from recombinant unlabeled NTD observed via phase contrast microscopy (C) Staining with Thioflavin T shows amyloid-like structures in cMLCK NTD condensates. (D) FRAP experiments with isolated cMLCK A647-NTD condensates. Data represent mean ± s.d. for N=4. Data in (A) were analyzed with a nested student's t-test after log transformation.

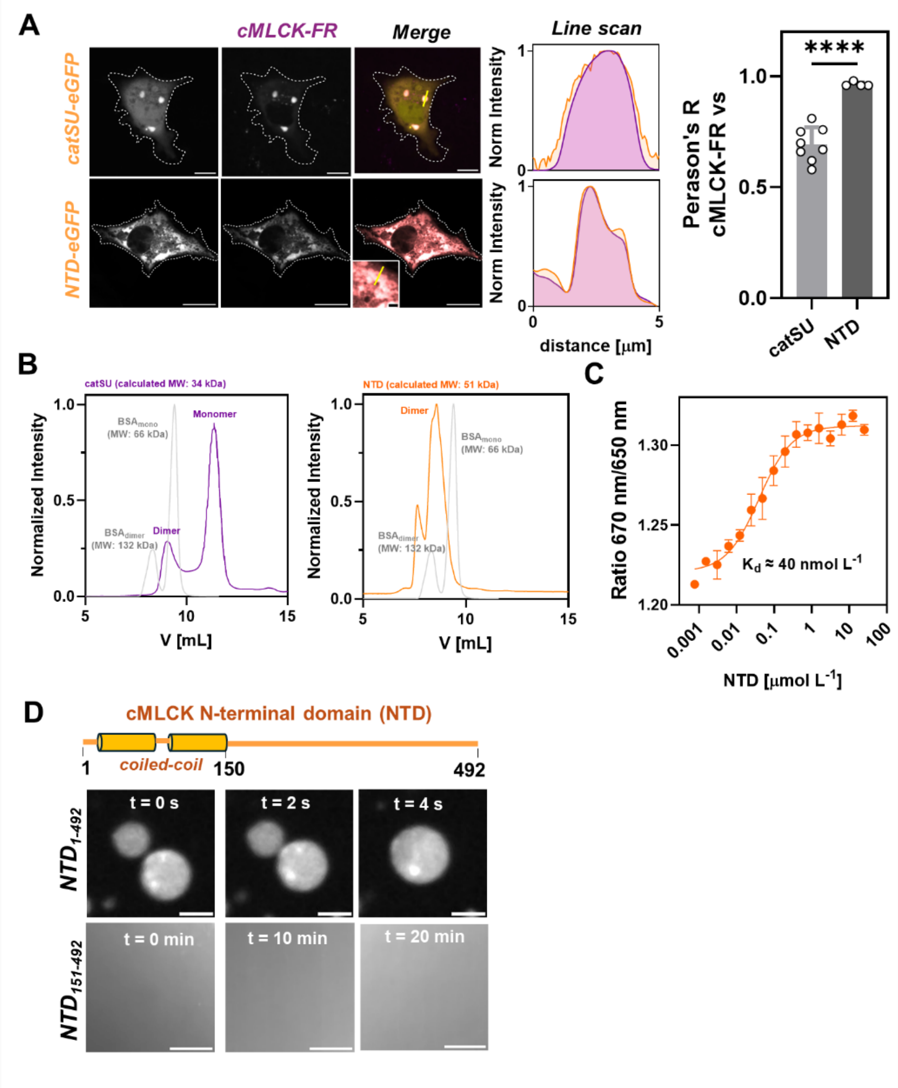


**Fig. S6. Intermolecular interactions stabilize cMLCK biomolecular condensates.** (A) Confocal image of live COS-1 cells co-expressing of catSU-eGFP (top) or NTD-eGFP with cMLCK-FusionRed (cMLCK-FR). Note that both catSU-eGFP and NTD-eGFP co-localize with cMLCK-FusionRed condensates. Pearson correlation coefficient analysis is shown on the right. Data are mean ± s.d. for n=4-8 cells. (B) Analytical gel filtration chromatograms for recombinant catSU (left, purple) and NTD (right, orange). Isolated bovine serime albumine (gray) was loaded as a control. catSU exists ina monomer-dimer equilibrium; whereas NTD is predomiantly a dimer in solution. (C) Microscale Thermophoresis analysis for the dimerization of NTD. Data represent mean ± s.d. for n=3 technical repeats. (D) Phase separation experiments of NTD in the presence ($NTD_{1\text{-}492}$) and in the absence of the predicted coiled-coil domain ($NTD_{151\text{-}492}$). Please note that $NTD_{151\text{-}492}$ does not phase separate into condensates.

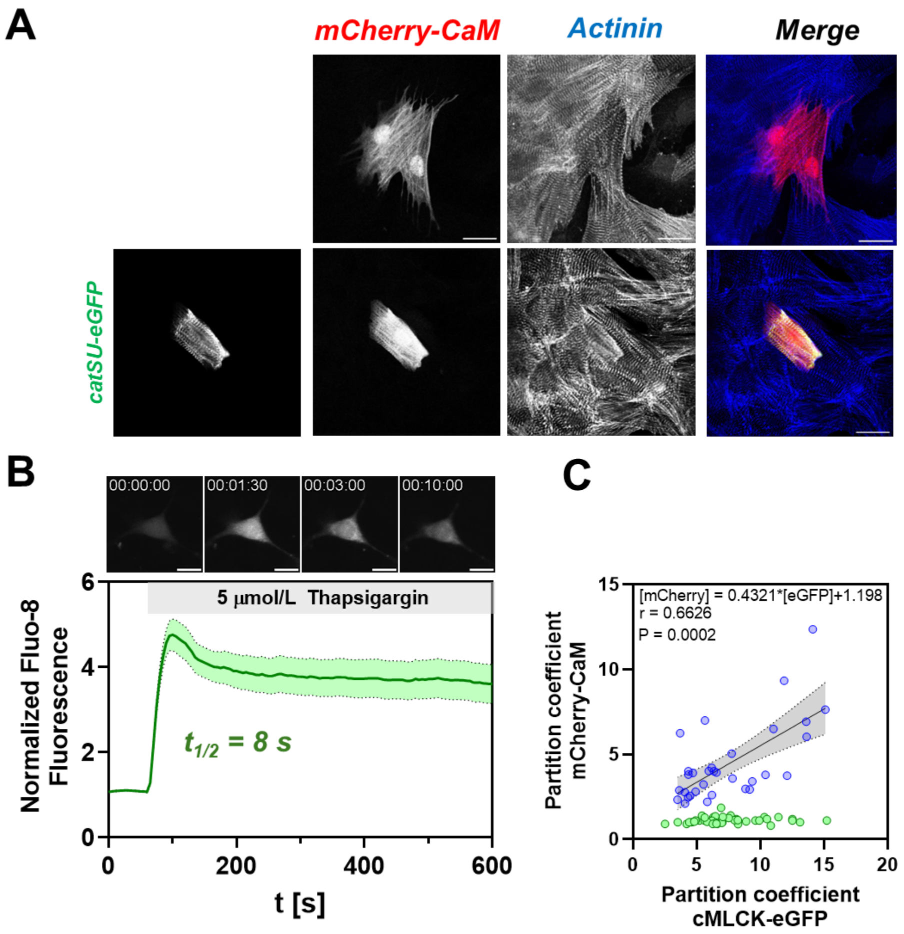


**Fig. S7. Calcium-dependent recruitment of calmodulin into cMLCK condensates.** (A) Confocal images of NRCs expressing mCherry-calmodulin (top) or co-expressing mCherry-calmodulin and the eGFP-tagges catalytic subunit (catSU) of cMLCK. (B) Calcium imaging of COS-1 cells using Fluo-8 during the Thapsigargin treatment protocol. Data represent mean ± s.e.m. for n=12 cells. (C) Plot of mCherry-CaM vs cMLKC-eGFP partition coefficient in COS-1 cells before (green) and after Thapsigargin treatment (blue). Linear regression to the data is shown as black continuous line. Grey shaded area indicates 95% confidence interval.

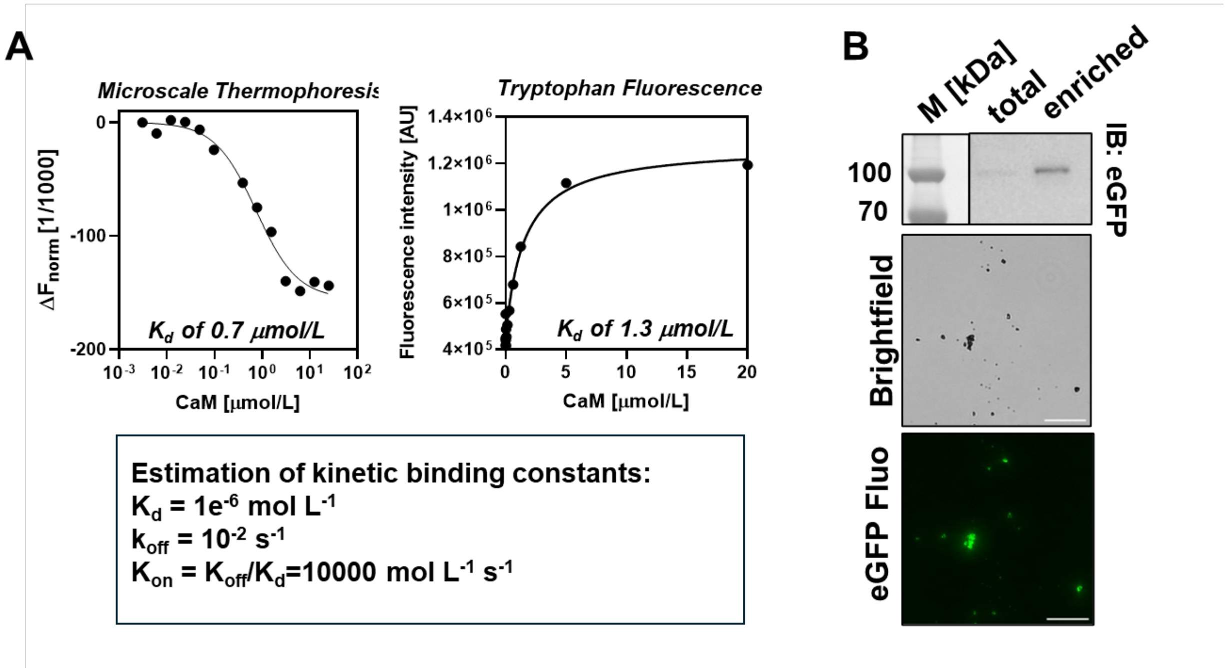


**Fig. S8. Binding affinity of calmodulin for cMLCK and purification of cMLCK condensates.** (A) Microscale Thermophoresis (left) and tryptophan fluorescence (right) binding isotherm for calmodulin binding to cMLCK. Data points were fitted to a single binding site model, extracting the steady state dissociation constant $K_d$. The box below shows the estimated kinetic constants of calmodulin binding to cMLCK. (B) Immuno-blot, and brightfield and fluorescence image of isolated cMLCK-eGFP condensates. Scale bar represents 20 μm.

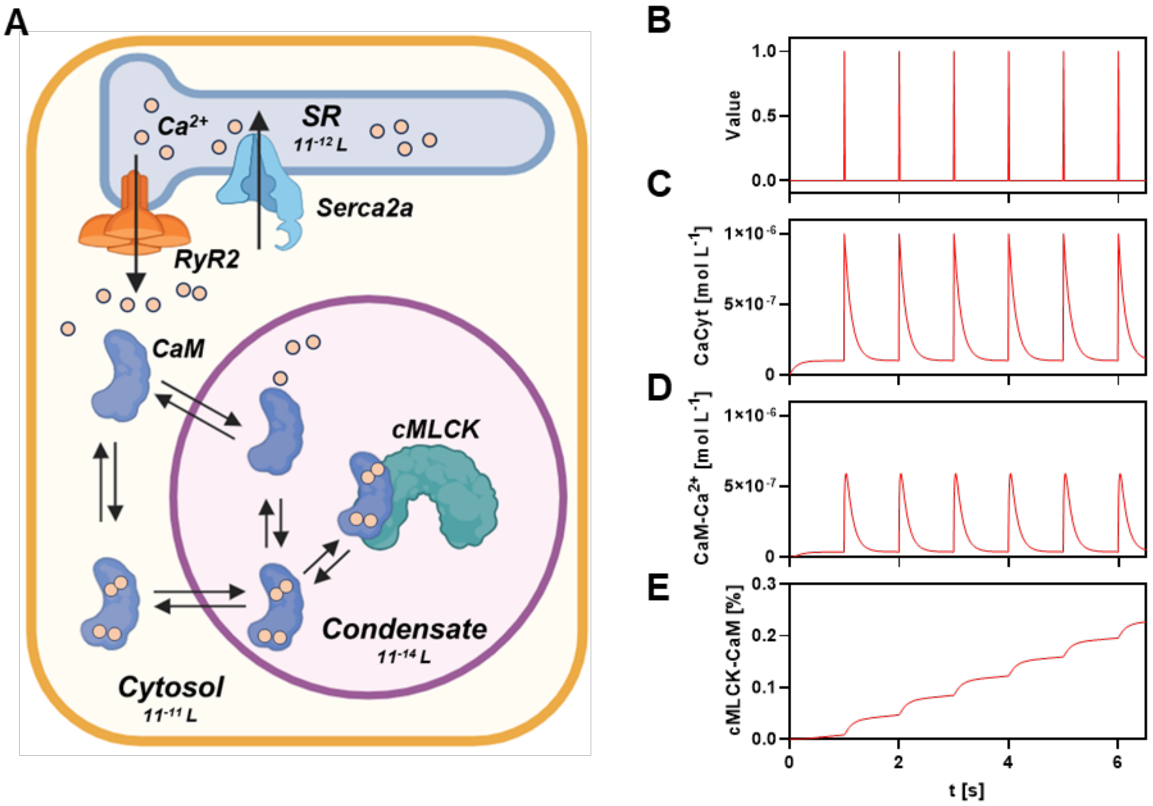


**Fig. S9. Cellular model of cMLCK activation.** (A) Cartoon representation of the biochemical model. (B) Activation pulse sequence simulating action potential. (C) Cytosolic calcium transient. (D) Concentration of $Ca^{2+}$-bound calmodulin. (E) Fraction of $Ca^{2+}$/calmodulin-activated cMLCK in condensates.

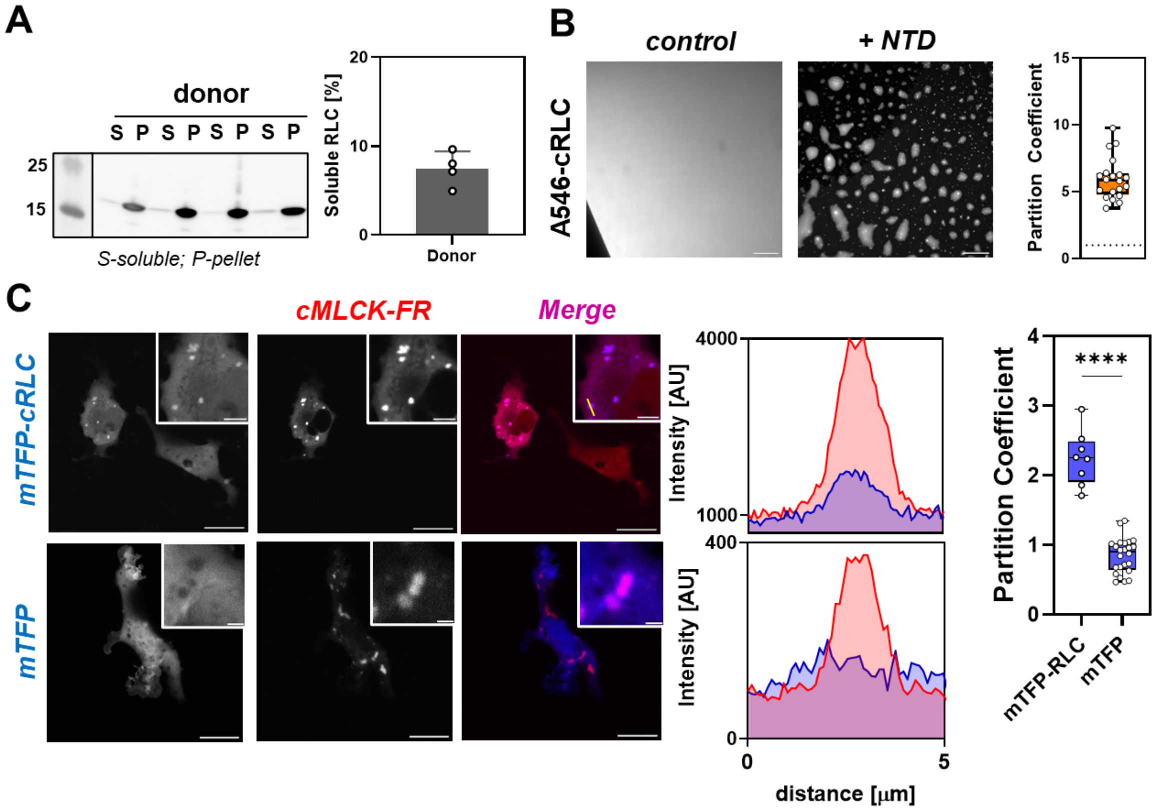


**Fig. S10. cMLCK condensates enrich cRLC.** (A) Fractionation of human ventricular tissue from organ donors and patients with heart failure into detergent-extracted soluble (S) and insoluble fraction (P) analyzed by immune-blotting against cRLC. Data represent mean ± s.d. (N=4 per group) and were analyzed using a one-way ANOVA followed by Tukey's post-hoc test. (B) Confocal image of Alexa 546 (A546) labeled cRLC alone (left) or in the presence of cMLKC NTD (right). The extracted partition coefficient is shown on the right (N=2, n=20 condensates). (C) Confocal image of COS-1 cell expressing FusionRed labeled cMLCK (cMLCK-FR) and mTFP-labled cRLC (top) or mTFP alone (bottom). Representative line scans are shown next to the confocal images. The comparison of partition coefficients of mTFP-cRLC and mTFP into cMLCK-R condensates are shown on the right (pooled data from n = 3 cells per condition). (D) Urea-glycerol gel of cRLC before and after preparative phosphorylation of serine 15 with recombinant cMLCK.

**Table S1. Primary and secondary antibodies used in this study.**

| **Primary Antibodies** | | |
|---|---|---|
| **Antigen** | **Supplier, catalogue number, host species** | **Application, dilution** |
| MYLK3 (cMLCK) | Proteintech, 21527-1-AP, rabbit polyclonal | IHC, 1:100<br>WB, 1:2000 |
| | Proteintech, 83673-3-RR, recombinant rabbit monoclonal | IHC, 1:100<br>WB, 1:2000 |
| MLRV (cRLC) | ABCAM, ab92721, rabbit monoclonal | WB, 1:2000 |
| Phospho MLRV (phospho cRLC) | Proteintech, AF861B, rabbit polyclonal | WB, 1:2000 |
| MYH7 (MHC) | DSHB, A4.1025, mouse monoclonal | IHC, 1:10 |
| TPM1 (cTm) | ABCAM, ab7785, mouse monoclonal | WB, 1:2000 |
| AT2A2 (SERCA2A) | Santa Cruz Biotechnologies, sc-376235, mouse monoclonal | WB, 1:1000<br>IHC, 1:100 |
| RYR2 | Invitrogen, MA3-916, mouse monoclonal | IHC, 1:100 |
| p62/SQSTM1 | Proteintech, 661B4, mouse monoclonal | IHC, 1:100 |
| GFP | ABCAM, ab1213, mouse monoclonal | IHC, 1:100 |
| ACTN (Actinin) | Sigma, clone EA-53, mouse monoclonal | IHC, 1:100 |
| **Secondary Antibodies** | | |
| Goat anti-rabbit-HRP | Invitrogen, 31460, polyclonal | WB, 1:2000 |
| Goat anti-mouse-HRP | Cell Signalling, 7076S, polyclonal | WB, 1:2000 |
| Goat anti-mouse Alexa 488 | ABCAM, ab150113, polyclonal | IHC, 1:100 |
| Goat anti-rabbit Alexa 594 | ABCAM, ab150080, polyclonal | IHC, 1:100 |
| Goat anti-mouse Alexa 594 | ABCAM, ab1500116, polyclonal | IHC, 1:100 |
| Goat anti-rabbit Alexa 488 | ABCAM, ab150077, polyclonal | IHC, 1:100 |

IHC – immunohistochemistry; WB – Western-blot

**Table S2. Plasmids used in this study.**

| Name | Vector | Insert | Comments |
|---|---|---|---|
| pET11a_NTD | pET3a | MYLK3_HUMAN (aa1-492) | N-terminal His-tag and TEV protease site |
| pET11a | pET11a | eGFP fused to MYLK3_HUMAN (aa1-492) | N-terminal His-tag |
| pET3a_NTDaa151-492 | pET3a | MYLK3_HUMAN (aa151-492) | N-terminal His-tag and TEV protease site |
| pET3a_Calmodulin | pET3a | CALM1_HUMAN | IN-terminal His-tag and TEV protease site |
| pET15b_cRLC | pET15b | MRLV_HUMAN (aa1-165) | N-terminal His-tag and TEV protease site |
| pET15b_NDcRLC | pET15b | MRLV_HUMAN (aa16-165) | N-terminal His-tag and TEV protease site |
| pET15b_cTnC | pET15b | TNNC1_HUMAN (aa1-161) | N-terminal His-tag and TEV protease site |
| pcDNA3.1(+)-C-eGFP | pcDNA3.1(+) | eGFP | |
| MYLK3_FL_pcDNA3.1(+)-C-eGFP | pcDNA3.1(+) | MYLK3_HUMAN (aa1-819) fused to eGFP | |
| MYLK3_N_pcDNA3.1(+)-C-eGFP | pcDNA3.1(+) | MYLK3_HUMAN (aa1-492) fused to eGFP | |
| MYLK3_catSU_pcDNA3.1(+)-C-eGFP | pcDNA3.1(+) | MYLK3_HUMAN (aa492-819) fused to eGFP | |
| mCherry-CaM_pcDNA3.1(+) | pcDNA3.1(+) | mCherry fused to CALM1_HUMAN | |
| cMLCK_FusionRed_pcDNA3.1(+) | pcDNA3.1(+) | MYLK3_HUMAN (aa1-819) fused to FusionRed | |
| mTFP_pcDNA3.1(+) | pcDNA3.1(+) | mTFP | |
| mTFP_cRLC_pcDNA3.1(+) | pcDNA3.1(+) | mTFP fused to MRLV_HUMAN (aa1-165) | |
